# Multiple interactions mediate the localization of BLTP2 at ER-PM contacts to control plasma membrane dynamics

**DOI:** 10.1101/2025.02.07.637094

**Authors:** Anbang Dai, Peng Xu, Chase Amos, Kenshiro Fujise, Yumei Wu, Han Yang, Julia N. Eisen, Andrés Guillén-Samander, Pietro De Camilli

**Author notes:** Correspondence (P.D.C).

## Abstract

BLTP2/KIAA0100, a bridge-like lipid transfer protein, was reported to localize at contacts of the endoplasmic reticulum (ER) with either the plasma membrane (PM) or recycling tubular endosomes depending on the cell type. Our findings suggest that mediating bulk lipid transport between the ER and the PM is a key function of this protein as BLTP2 tethers the ER to tubular endosomes only after they become continuous with the PM and that it also tethers the ER to macropinosomes in the process of fusing with the PM. We further identify interactions underlying binding of BLTP2 to the PM, including phosphoinositides, the adaptor proteins FAM102A and FAM102B, and also N-BAR domain proteins at membrane-connected tubules. The absence of BLTP2 results in the accumulation of intracellular vacuoles, many of which are connected to the plasma membrane, pointing to a role of the lipid transport function of BLTP2 in the control of PM dynamics.

## Introduction

Cellular life implies continuous intracellular fluxes of lipids. This is achieved both by vesicular transport and by protein mediated lipid transport. Until few years ago, most lipid transport proteins were thought to function via a shuttle mechanism in which lipid binding proteins extract lipids from a membrane and transport them piecemeal to another membrane, often in a counter-transport reaction whereby two different lipids are exchanged between the two membranes^1–3^. Recently, evidence for another mechanism of lipid transport between membranes has been reported, mediated by proteins that directly bridge two different membranes and contain a hydrophobic groove spanning their entire length along which lipids can slide from one membrane to another^4–7^. These proteins, collectively referred to as bridge-like lipid transfer proteins (BLTPs), are optimally suited for fast high-capacity lipid transport. Functions assigned to these proteins include growth of new membranes, expansion of organelles not connected to other membranes by vesicular transport, membrane repair and roles in the rapid remodeling of the lipid composition of membranes^8^.

BLTP2/KIAA0100 is one such protein with orthologues in all eukaryotic species^4,5,7^. It is a large protein predicted to have a rod-like structure with an N-terminal transmembrane helix that anchors it in the ER. In mammals its absence results in embryonic lethality, in flies where it was formerly referred to as “hobbit”, it is essential for development and synaptogenesis^9,10^ and in plants for root growth and cytokinesis^11^. Genetic studies of human BLTP2 and of its orthologues in yeast (Fmp27 and Ypr117w) and in worms (F31C3.3), have suggested its role in the adaptation of life to lower temperature^7,12^, most likely via their property to rapidly modify the membrane lipidome to ensure normal membrane fluidity at reduced temperature (known as homeoviscous adaptation). A similar function has been assigned to BLTP1 in yeast (Csf1)^7,13,14^ and worms (LPD-3)^15,16^. Concerning its site(s) of action within cells, studies in yeast, Drosophila cells and a human cancer cell line reported its concentration at ER-Plasma Membrane (PM) contacts^7,12,17^, while another study of mammalian cell lines reported its localization at contacts between the ER and tubular recycling endosomes^18^, rather than at ER-PM contacts. A localization of yeast Fmp27 and Ypr117w at some ER-mitochondria contacts was also reported^7^.

The goal of this study was to advance our understanding of the localization and function of BLTP2. We consistently found BLTP2 at contacts between the ER and the PM, although in different patterns depending on the cell line. In cells where BLTP2 was localized at contacts of the ER with apparently internal membranes, such as tubular endosomes or macropinosomes, we found that such membranes were continuous with, or in the process of becoming continuous with, the PM. We also identified interactions responsible for these localizations, including the binding of BLTP2 to two PM adaptors, FAM102A and FAM102B. Finally, we found that cells lacking BLTP2 have striking alterations of their internal structure with the abundant presence of intracellular vacuoles positive for PM markers and open to the cell surface. These alterations reveal a function of BLTP2 in controlling the dynamics of the PM which may result from a defect in BLTP2 dependent lipid transport to this membrane.

## Results

### BLTP2 localize at contact of the ER with the PM, including deep PM invaginations continuous with tubular recycling endosomes

To investigate the cellular localization of BLTP2 in human cells, we first expressed, transiently or constitutively through a lentivirus, BLTP2 constructs with an internal fluorescent tag (BLTP2^Halo or BLTP2^EGFP) (Figure 1A) in different cell types (U2OS cells, MDA-MB-231 cells, HeLaM cells and COS-7 cells). The tag was inserted in a loop emerging from its predicted rod-like core at a site not predicted to affect BLTP2 folding based on AlphaFold3^19^ (Figure 1B), as the C-terminal region of BLTP2 had been implicated in tethering ER-anchored BLTP2 to other membranes in Drosophila cells^17^. Thus, we wished to avoid perturbation of this interaction, which appears to be critical for BLTP2 function.

**Figure 1.**
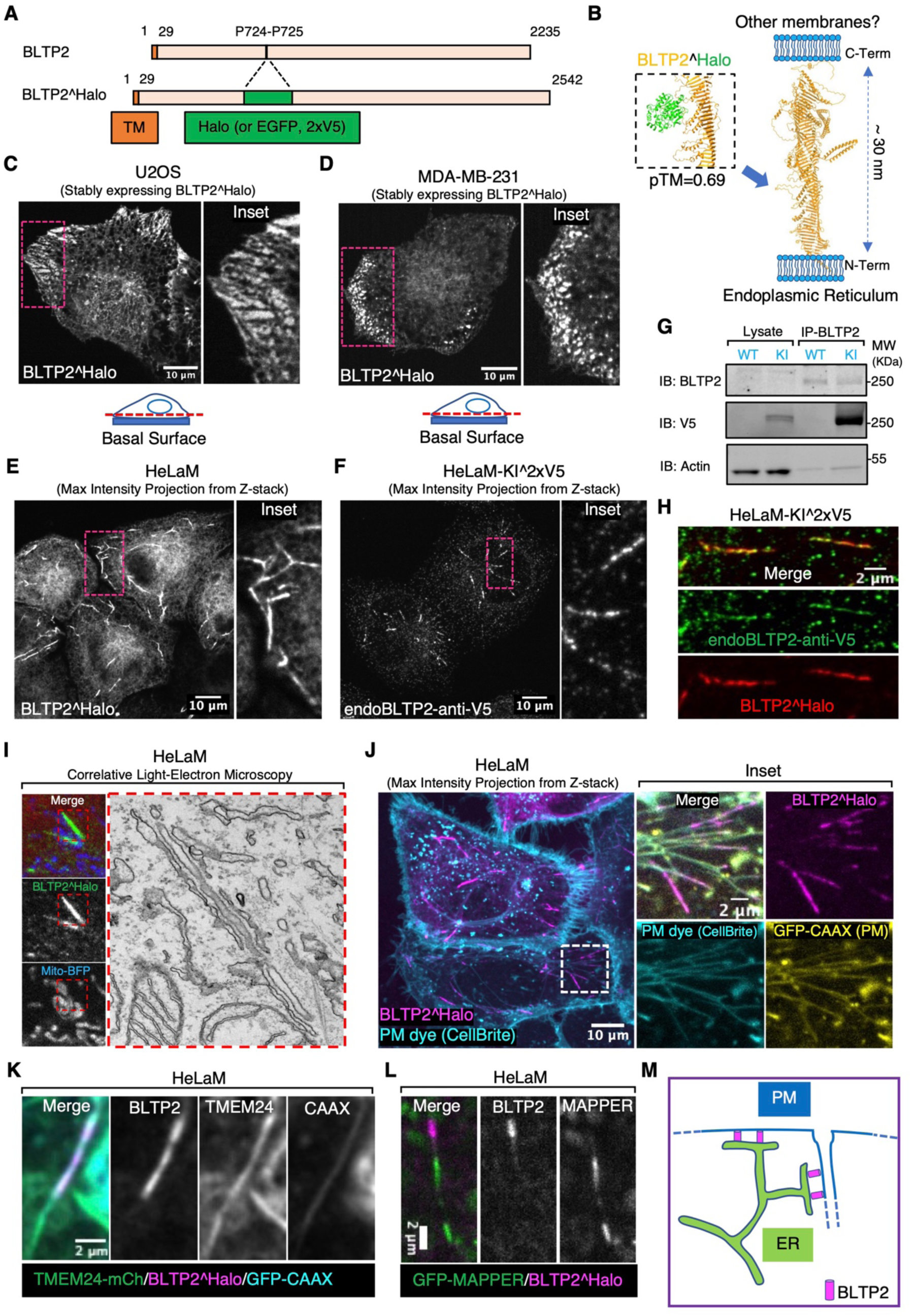
BLTP2 is enriched at ER contacts with the PM and PM-connected tubular internal membranes. (A) Domain diagram of human BLTP2 and of the internally tagged (Halo, EGFP or 2xV5 epitopes) BLTP2. TM indicates the transmembrane region of BLTP2. (B) Schematic model of the arrangement of BLTP2 at contacts of the ER with other membranes. The arrow indicates the site where tags (Halo, EGFP or 2xV5 epitopes) were inserted. Structures are predicted using AlphaFold3. (C and D) U2OS cells (C) and MDA-MB-231 cells (D) stably expressing BLTP2^Halo show enrichment of this protein at ER-PM contacts near the edge of the cell. An optical section close to the basal surface (see dashed red line in the cartoon) is shown. (E) HeLaM cells stably expressing BLTP2^Halo show enrichment of the protein at tubular structures. (F) Localization of endogenous BLTP2 in gene edited HeLaM cells where the 2xV5 epitope was inserted in the coding sequence of BLTP2. Anti-V5 immunofluorescence reveals enrichment of BLTP2 on tubular structures. (G) Validation of the endogenous tagging of BLTP2 by western blotting. Anti-BLTP2 immuno-precipitation coenriched a V5 immunoreactive band. (H) Endogenous BLTP2 (V5 immunoreactivity) co-localizes with exogenous BLTP2^Halo on the same tubular structures. (I) Correlative Light-Electron Microscopy revealed that a BLTP2-positive tubular structure represents a tubular membrane surrounded by ER. (J) BLTP2^Halo localizes at the distal portion of tubular structures that are positive for the PM marker GFP-CAAX and are labeled by the membrane impermeant PM dye CellBrite. (K) TMEM24-mCherry, another ER-PM contact protein, is also present on BLTP2-positive tubular structures but only partially colocalizes with BLTP2^Halo. (L) GFP-MAPPER, an artificial ER-PM tethering protein, is also present on BLTP2-positive tubular structures but does not overlap with BLTP2^Halo. (M) Schematic drawing of BLTP2 localization at contacts of the ER with both the “outer” PM and the distal portions of PM-connected tubular structures.

Consistent with BLTP2 being a resident integral membrane protein of the ER^7,9,17,18^, tagged BLTP2 produced a diffuse ER fluorescence, as revealed by the co-expression of the ER marker RFP (or iRFP)-Sec61β (Figures S1A, S1B and S1D). However, in addition to this diffuse ER fluorescence, focal accumulations of BLTP2^Halo were observed with cell line specific differences. In U2OS and MDA-MB-231 cells, BLTP2 positive-patches with the typical appearance of ER-PM contacts were visible, primarily at the basal surface of these cells and concentrated at the leading edge in migrating cells (Figures 1C, S1A, 1D and S1B). In HeLaM cells such patches were not visible and BLTP2^Halo was instead concentrated at the distal portion of tubular internal structures that radiated from the center to the cell periphery (Figure 1E). This localization reflected the native localization of BLTP2 in these cells as it overlapped with anti-V5 immunofluorescence of gene edited HeLaM cells harboring a twin-V5 epitope tag in endogenous BLTP2 (at the same internal site used to tag with Halo exogenous BLTP2) (Figures 1F-1H). Similar BLTP2-positive tubules were only occasionally observed in MDA-MB-231 cells (Figure S1C). In COS-7 cells, BLTP2 hot spots either in the form of tubules (as in HeLaM cells) or of small patches at the cell periphery were only infrequently observed (Figures S1D and S1E).

The selective concentration of both exogenous and endogenous BLTP2 on tubular structures positive for Rab8 and Rab10 in HeLa cells was previously reported and thought to reflect contacts between the ER and tubular recycling endosomes^18^. In agreement with this previous study, we detected the presence of Rab8 and Rab10 on the BLTP2-positive tubules in HeLaM cells (Figures S4A and 2A). We confirmed by correlative-light electron microscopy (CLEM) that BLTP2-positive linear structures corresponded to contacts between the ER and membrane tubules (Figure 1I). We found, however, that at least the great majority of BLTP2-positive tubular structures were positive for PM markers, such as GFP-CAAX^20^ or Lyn11-RFP^21^ (Figures 1J, S2A and S2B). Accordingly, these tubular structures became labeled with the non-permeable membrane dye CellBrite® steady 650 after few minutes incubation at 37 ℃ (Figures 1J, 2B and S2A) or after 15 mins incubation at 4 ℃ (Figure S2C). Moreover, addition of antibodies directed against the ectodomain of the Major Histocompatibility Complex Class I (MHC-I) complex to live HeLaM cells expressing BLTP2^EGFP resulted not only in a diffuse fluorescence of cell surface, where the bulk of MHC-I is localized, but also in the labeling of the BLTP2-positive tubules (Figure S4B), in agreement with the known presence of these complexes in tubular endosomes^22–24^. Other ER-PM tethers, such as TMEM24-mCherry^25^ or GFP-MAPPER^26^ (a synthetic ER-PM tether), were also found at the tubular structures (Figures 1K and 1L). However, these other proteins only partially co-localized with BLTP2 on the tubules and were additionally concentrated at other ER-PM contacts, such as those on the basal surface of the cell, where BLTP2 was not observed. We conclude that while in HeLaM cells BLTP2 localizes at contacts with tubular internal membranes with the properties of tubular recycling endosomes, such membranes are continuous with the PM and thus represent extensions of the PM. These tubules are likely the same as the recently described Rab10 positive tubules open to the PM^27^, which were referred to as PM invaginations rather than as recycling endosomes. The presence of BLTP2 only on the most distal (peripheral) portion of the tubules indicate that despite their continuity with the PM, a heterogeneity is maintained along them with a progressive transition from PM-like properties in their peripheral portions to *bona fide* endosomal properties towards the cell center.

### Tubular recycling endosomes acquire BLTP2-positive contacts as they fuse with the PM

The CAAX-positive tubules surrounded by BLTP2 in their distal portion were Rab10-dependent as they were no longer observed in cells expressing dominant negative Rab10 mutant (Rab10-T23N) (Figures 2C-2G). Formation of Rab10-dependent tubules, in turn, are known to be driven by microtubules (MTs) based motor proteins, KIF13A and KIF13B^28^. Accordingly, they were disrupted by depolymerization of microtubules using nocodazole as described^28^, but reformed starting from the central region of the cell after washout the drug (Figure 2H). This property offered us the opportunity to monitor the appearance of BLTP2 signal upon their re-growth. We found that BLTP2 was recruited at their tips only when they reached the cell periphery and became continuous with the cell surface (Figures 2I and 2J; Video S1).

**Figure 2.**
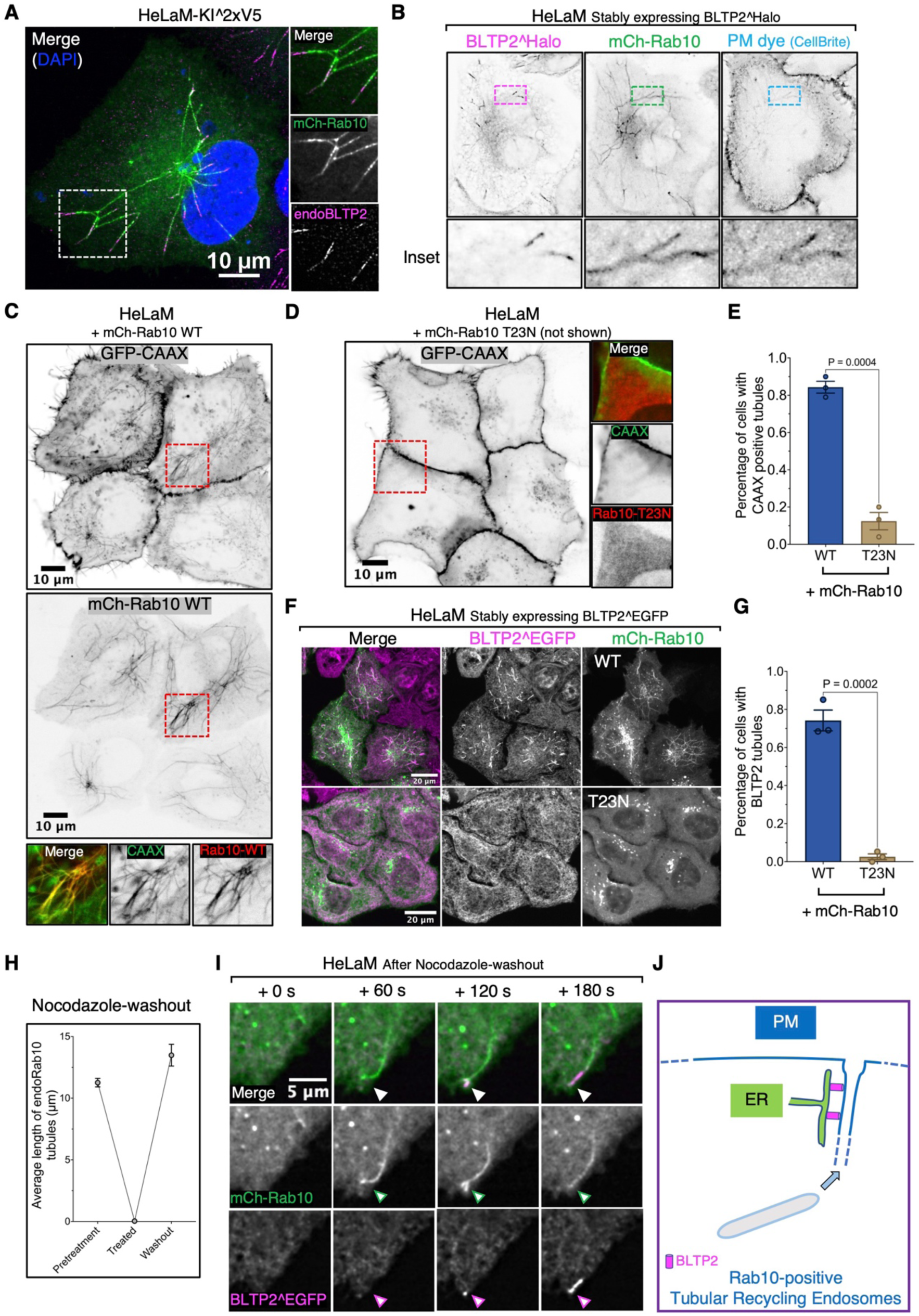
BLTP2-positive tubular internal membranes are Rab10-dependent tubular recycling endosomes continuous with the PM. (A) Endogenous BLTP2 localize at the tip of mCherry-Rab10-positive tubular endosomes in HeLaM cells. (B) BLTP2^Halo- and mCherry-Rab10-positive tubular endosomes in HeLaM cells are connected with the PM as revealed by labeling with the membrane impermeant PM dye CellBrite. (C) mCherry-Rab10-positive tubular endosomes in HeLaM cells are also labeled with the PM marker GFP-CAAX. (D and E) Absence of GFP-CAAX positive tubules in HeLaM cells expressing dominant negative Rab10 (mCh-Rab10 T23N). Fluorescence image in (D) and quantification in (E). Two tailed t-test. Mean ±SEM. n=3 independent experiments. 48 cells for WT and 57 cells for the T23N Mutant. (F and G) Expression of dominant negative Rab10 (mCh-Rab10 T23N) abolished BLTP2^EGFP-positive tubules. Fluorescence images in (F) and quantification in (G). Two tailed t-test. Mean ±SEM. n=3 independent experiments. 127 cells for WT Rab10 and121 cells for the T23N mutant. (H) Tubular endosomes positive for endogenous Rab10 immunoreactivity disappear after nocodazole treatment for two hours but are restored after washing out the drug. n=3 independent experiments. Pretreatment: 269 Rab10 tubules from 28 cells; treated: zero tubules observed in the 46 cells examined; washout: 353 Rab10 tubules from 26 cells. (I) Live-imaging of a HeLaM cells after nocodazole washout shows the recovery of a Rab10-positive tubular endosome and the recruitment of BLTP2^EGFP after the tubule reaches the PM and fuses with it. (J) Schematic drawing of BLTP2 recruitment and localization at a Rab10-positive tubular endosome that is continuous with the PM.

### Recruitment of BLTP2 to PM-connected tubular recycling endosomes requires PI4P generated by PI4KIIIα in their membrane

The results reported above indicate that a shared feature of the ER contact sites populated by BLTP2 is to be with the PM, in spite of cell-specific differences in their localization within this membrane. We next investigated PM determinant responsible for these localizations.

Many proteins that function as ER-PM tethers bind PI4P and PI(4,5)P_2_, two phosphoinositides concentrated in the PM^29^. As stated above, BLTP2 comprises a C-terminal region which contains numerous basic amino acids (a.a.) whose deletion in Drosophila (a.a.2219-2300) abolishes its localization at ER-PM contacts and for its physiological function^17^. Accordingly, we tested the effect of the deletion of the C-terminal region in human BLTP2 and found that the removal of the last 59 a.a. (a.a.2177-2235), which has an overall basic charge, was sufficient to reduce its concentration not only at the “outer” PM (Figures S3A and S3B), but also at the surface-connected membrane tubules (Figure S3C) where both PI4P and PI(4,5)P_2_ are concentrated as revealed by expression of iRFP-P4M^30^ and GFP-PH_PLCδ1_^31,32^ respectively (Figure 3A). To confirm the role of these two phosphoinositides in BLTP2 localization, we used the rapamycin-dependent FKBP-FRB heterodimerization system^21,33^ to acutely recruit phosphoinositide phosphatases to the PM to deplete them (Figure 3B). We co-expressed a “bait” construct comprising the PM targeting sequence of Lyn11 followed by the FRB domain (Lyn11-CFP-FRB) and a “prey” construct comprising the FKBP domain followed either by the 4-phosphatase domain of Sac1 (to dephosphorylate PI4P to PI) (mRFP-FKBP-Sac1)^34^, or the 5-phosphatase domain of INPP5E (to dephosphorylate PI(4,5)P_2_ to PI4P) (mRFP-FKBP-INPP5E)^35^. Upon rapamycin addition to induce heterodimerization, robust recruitment of either prey constructs from the cytosol to the tubules was observed. However, only after the recruitment of mRFP-FKBP-Sac1 BLTP2 disassociated from the tubules, revealing a critical role of PI4P (Figure 3C).

**Figure 3.**
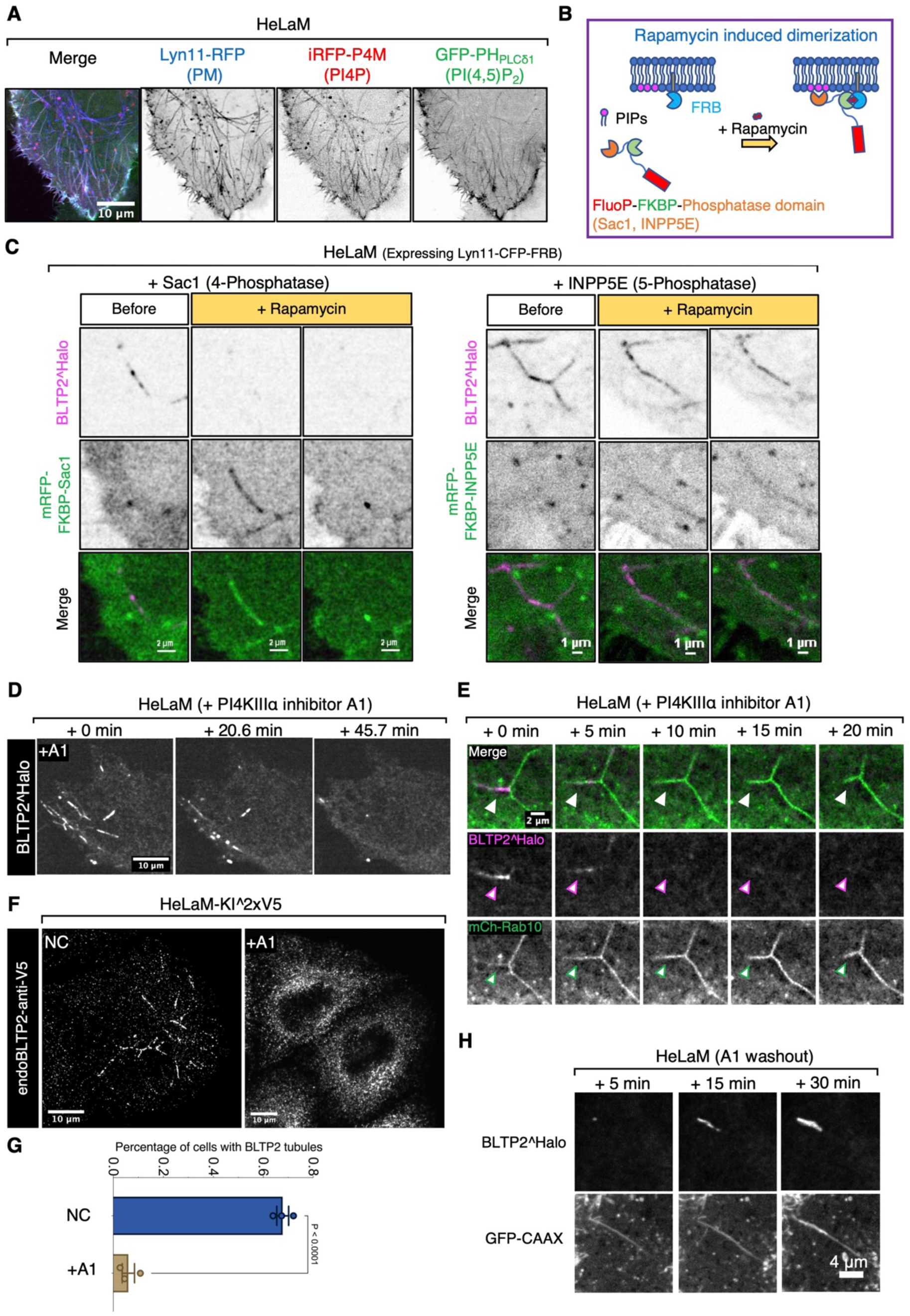
PI4P regulates BLTP2-dependent contacts of the ER with tubular endosomes. (A) PM-connected tubular endosomes in HeLaM cells are positive for PI4P (labeled by iRFP-P4M) and PI(4,5)P_2_ (labeled by GFP-PH_PLCδ1_) markers. (B) Design of the rapamycin-dependent dimerization assay to recruit the 4-phosphatase domain of Sac1 (target PI4P) or the 5-phopshatase domain of INPP5E (target PI(4,5)P_2_) to PM-connected tubular endosomes. (C) BLTP2^Halo disassociates from tubular endosomes after PI4P depletion on their membranes following the recruitment of RFP-FKBP-Sac1. In contrast, BLTP2^Halo show no clear change after PI(4,5)P_2_ depletion following the recruitment of RFP-FKBP-INPP5E. (D) BLTP2^Halo gradually disassociates from tubular endosomes after PI4KIIIα inhibition in response to addition of the A1 compound. (E) mCh-Rab10 tubular endosomes persist after BLTP2^Halo disassociation from them in response to A1 treatment. (F) Endogenous BLTP2 also undergoes disassociation from tubular endosomes in response to treatment with A1. (G) Cells in (F) are quantified for the presence of BLTP2 tubules. Two-tailed t-test. Mean ±SEM. n=3 independent experiments. Non treated (NC): 85 cells; A1 treated: 118 cells. (H) BLTP2^Halo re-associates with tubular endosomes after A1 washout.

PI4P localized at the PM is primarily synthesized by PI4KIIIα, the kinase encoded by the PI4KA gene^36^. Supporting the importance of the PM pool of PI4P in the binding of ER-anchored BLTP2 to the tubules, addition of the A1 compound, a specific PI4KIIIα inhibitor which in the absence of triggered PI(4,5)P_2_ hydrolysis, selectively decreases PI4P but not PI(4,5)P_2_ at the PM^37,38^, induced the dissociation of both BLTP2^Halo (Figure 3D) and endogenous BLTP2 from the tubules (Figures 3F and 3G) although the tubular network was not disrupted (Figures 3E and S5A; Video S2). This reaction was also reversible, as washout of the drug after 1 hour treatment resulted in the reformation of BLTP2 positive contacts on the tubules (Figures 3H and S5B; Video S3). We conclude that PI4P, but not PI(4,5)P_2_, is required for the tethering function of BLTP2 at these sites. While these results reveal the importance of PI4P in BLTP2 recruitment, PI4P is unlikely to represent the unique determinant for the concentration of BLTP2-dependent contacts at a specific PM sub-compartment. PI4P may function as an essential coreceptor. Thus, we considered the possible occurrence of protein receptors for BLTP2 in the PM.

### FAM102A and FAM102B are PM adaptors for BLTP2

A recently posted database of the yeast interactome^39^ (http://yeast-interactome.biochem.mpg.de:3838/interactome/) revealed that the two yeast orthologs of BLTP2, Fmp27 and Ypr117w are interactors of Ybl086c, with Fmp27 being the strongest interactor (Figures 4A and 4B). Interestingly, other interactors of Fmp27 are evolutionarily conserved proteins also implicated in lipid dynamics at ER-PM contacts (Ist2^40^ and Osh3^41^) or metabolism (Lro1^42^). Ybl086c, in turn, has two orthologues in humans, FAM102A and FAM102B, which are similar proteins comprising an N-terminal C2 domain and a C-terminal disordered region (Figures 4C and S6A). FAM102A, also named as EEIG1 (Early Estrogen induced gene 1), was identified as a RANK (Receptor Activator of Nuclear factor κB) ligand involved in the regulation of osteoclast formation and bone remodeling^43–45^. However, the cellular functions of both FAM102A and FAM102B are not known. As C2 domains can function as bilayer binding modules, we explored the possibility that these proteins could function as potential membrane adaptors for BLTP2 in mammalian cells.

**Figure 4.**
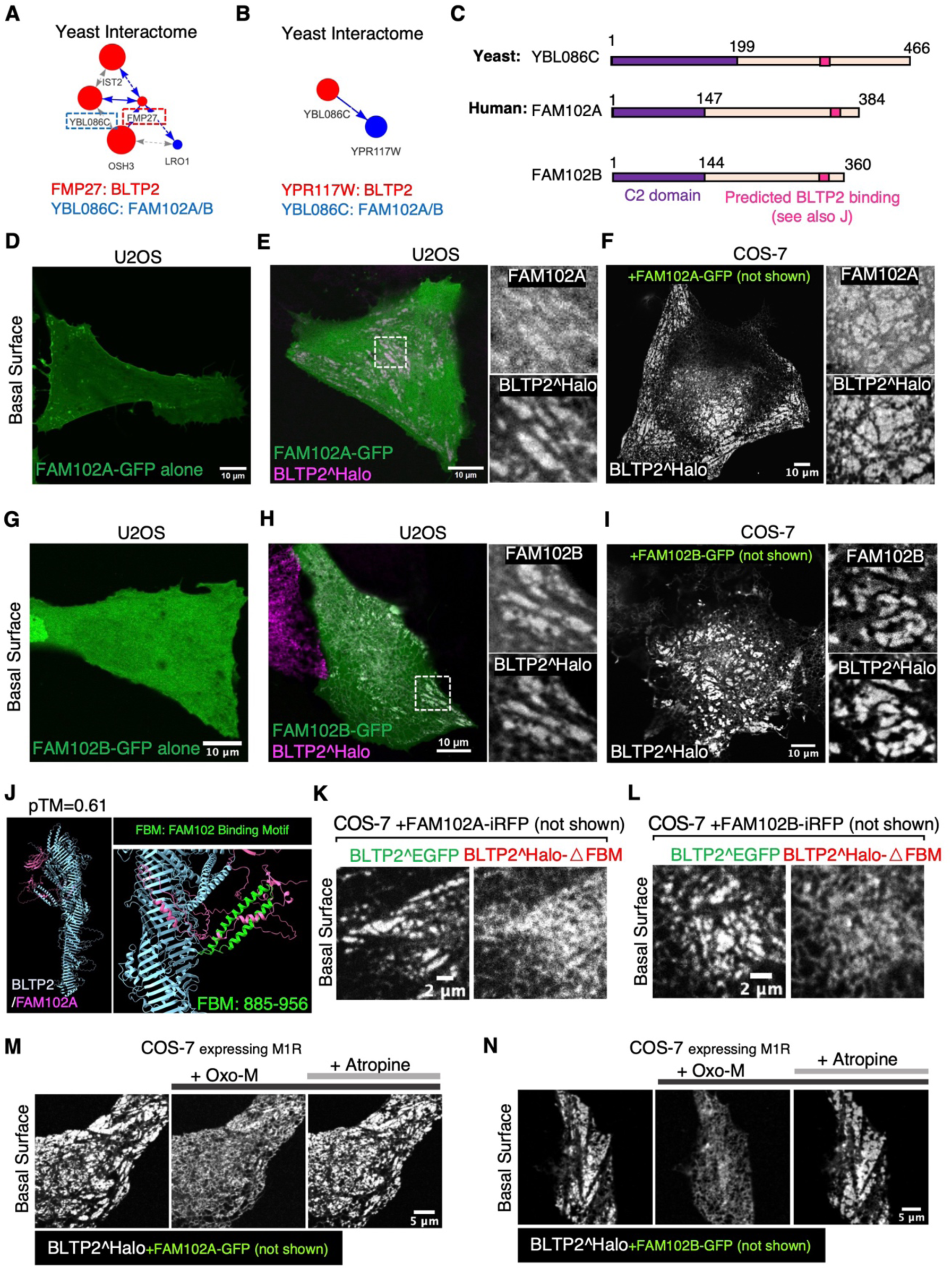
An interaction of BLTP2 with FAM102A and FAM102B enriches BLTP2 at ERPM contacts. (A and B) Interaction diagram exported from the Yeast Interactome Website (http://yeast-interactome.biochem.mpg.de:3838/interactome/) revealing protein interactions of the BLTP2 yeast orthologues Fmp27 (A, score=10) and Ypr117w (B, score=3) with Ybl086c, the ortholog of mammalian FAM102A/B. (C) Domain organization of Ybl086c with FAM102A/B comprising an N-terminal C2 domain and a C-terminal disordered region. The AlphaFold3-predicted BLTP binding site in FAM102A and B is shown in (J) (see also Figure S6A). (D) Solo expression of FAM102A-GFP shows localization at the PM in U2OS cells, with no focal accumulations as expected for ER-PM contact sites. (E) In U2OS cells co-expressing FAM102A-GFP and BLTP2^Halo, the two proteins colocalize at ER-PM contacts. (F) In COS-7 cells, where BLTP2 does not accumulate at ER-PM contacts when expressed alone, co-expression of FAM102A-GFP recruits BLTP2^Halo to ER-PM contacts. (G) Solo expression of FAM102B-GFP in U2OS cells results in its diffuse localization. (H) In U2OS cells co-expressing FAM102B-GFP and BLTP2^Halo, the two proteins colocalize at ER-PM contacts. (I) In COS-7 cells, where BLTP2 does not accumulate at ER-PM contacts when expressed alone, co-expression of FAM102B-GFP recruits BLTP2^Halo to ER-PM contacts. (J) AlphaFold3 predicts an interaction between the C-terminal region of FAM102A (magenta) and a two-helices hairpin (green) (Fam102 Binding Motif, FBM) projecting out of the BLTP2 rod-like core (blue). (K and L) COS-7 cells co-expressing BLTP2^EGFP and BLTP2^Halo-△FBM with FAM102A-iRFP (K) or FAM102B-iRFP (L), respectively, showing that BLTP2^Halo-△FBM is not co-enriched with WT BLTP2^EGFP at ER-PM contact sites, but remains diffuse throughout te ER. (M and N) In COS-7 cells co-expressing BLTP2^Halo, the M1R receptor and either FAM102A-GFP (M) or FAM102B-GFP (N), BLTP2^Halo disassociates from the ER-PM contact sites and redistributes to the entire ER in response to PI(4,5)P_2_ depletion by Oxo-M addition. BLTP2^Halo re-localizes to ER-PM contacts after addition of the Oxo-M antagonist atropine.

First, we examined their localization when expressed as C-terminally-tagged fusion proteins in U2OS cells or COS-7 cells. When expressed alone, both FAM102A-GFP and FAM102B-GFP had a diffuse PM and cytosolic localization (Figures 4D and 4G). However, when co-expressed with exogenous (and thus overexpressed) BLTP2^Halo, both FAM102A and FAM102B colocalized with BLTP2 at PM patches with the typical morphology of ER-PM contacts not only in U2OS cells (Figures 4E and 4H) and MDA-MB-231 cells (Figure S6B), but also in COS-7 cells (Figures 4F and 4I), where BLTP2^Halo, as mentioned above, does not exhibit an obvious concentration at ER-PM contacts (Figure S1D). These results suggest that FAM102A/B may help recruit BLTP2 to the ER-PM contacts, but they may be present in limiting concentration in COS-7 cells.

We then used AlphaFold3 to determine whether the FAM102 proteins may directly bind BLTP2 and found a high confidence (pTM=0.61) predicted interaction between a conserved sequence in the C-terminal region of FAM102A/B and an alpha-helix hairpin (hence called FBM for <u>F</u>AM <u>B</u>inding <u>M</u>otif) that projects out from the rod core of BLTP2 (Figures 4J and S6C). Supporting the physiological importance of this interface, a BLTP2 deletion construct lacking this motif (BLTP2^Halo-△FBM) was no longer recruited to PM patches by co-expression with either FAM102A or FAM102B in COS-7 cells, in contrast to full length (FL) BLTP2^EGFP (Figures 4K and 4L).

We found that FAM102-BLTP2 positive ER-PM contacts are regulated by PI(4,5)P_2_, as both FAM102 proteins and BLTP2 disassociated from the PM when PI(4,5)P_2_ was acutely depleted through phospholipase activation driven by the addition of Oxo-M to COS-7 cells also expressing the muscarinic receptor M1R^46^. These contacts were re-established after adding the muscarinic receptor antagonist atropine (Figures 4M and 4N; Videos S4 and S5).

Surprisingly, when we expressed FAM102A and FAM102B in HeLaM cells, we observed that FAM102B was selectively enriched on the tubular structures (Figure 5A), where it precisely colocalized with both exogenous and endogenous BLTP2 (Figures 5B and 5C), while FAM102A had a diffuse localization at the PM (Figure 5A). However, when BLTP2^Halo was co-expressed with FAM102A-GFP, the two proteins colocalized at typical ER-PM patches on the basal surface of the cell (Figure S6D), suggesting that the selective localization of BLTP2 on the tubule in the absence of FAM102A overexpression reflects the predominant expression of FAM102B in these cells.

**Figure 5.**
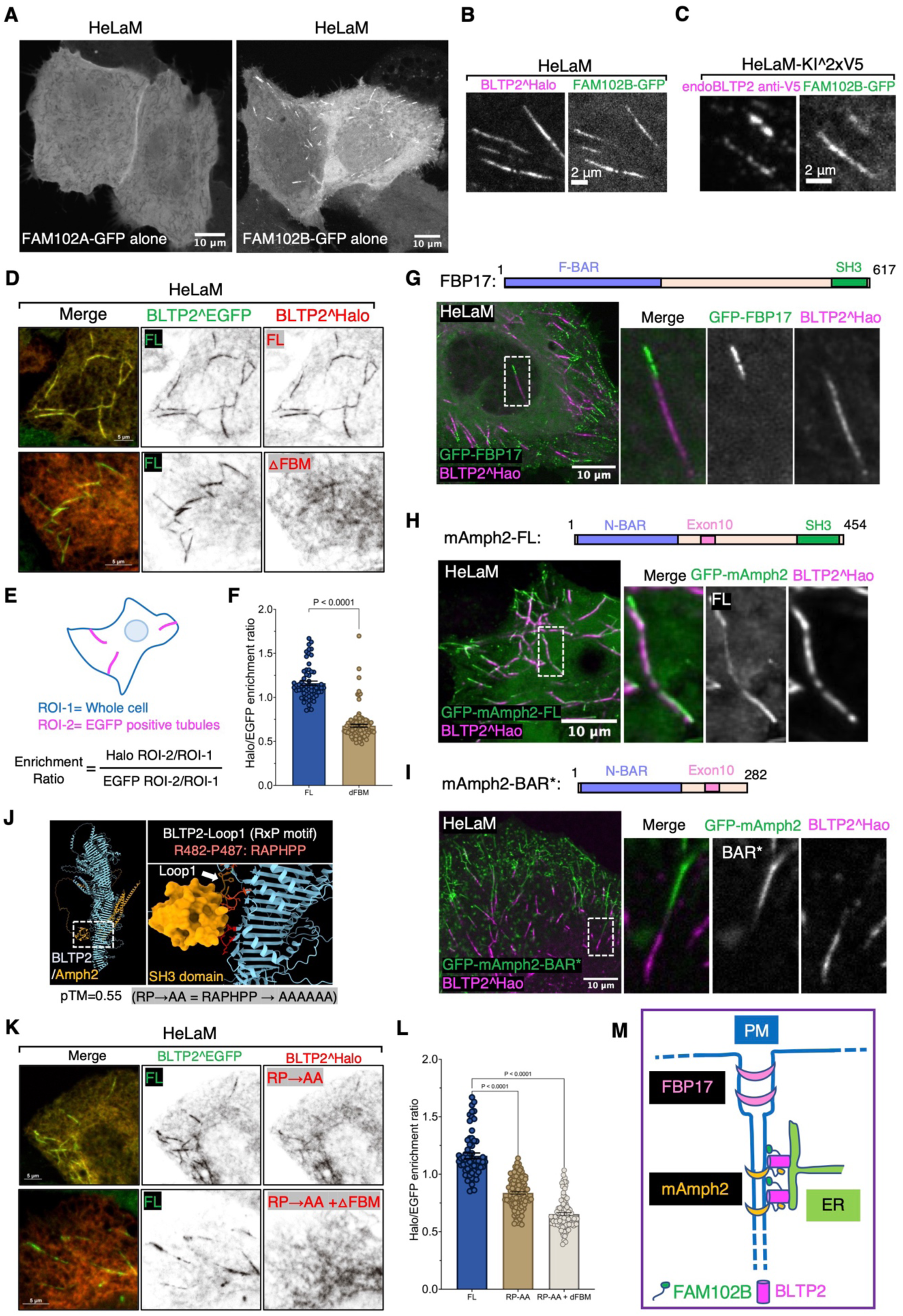
Interactions of BLTP2 with FAM102B and N-BAR domain proteins at PM-connected tubular endosomes in HeLaM cells. (A) Solo expression of FAM102A-GFP in HeLaM cells results in its diffuse localization, while solo expression of FAM102B-GFP results in its enrichment localization on tubular structures similar to BLTP2 localization. (B and C) FAM102B-GFP co-localizes with either BLTP2^Halo (B), or endogenous BLTP2 (C) on tubular structures in HeLaM cells. (D-F) Co-expression of BLTP2^EGFP FL (full length) with either BLTP2^Halo FL or BLTP2^Halo-△FBM, respectively, in HeLaM cells, showing that deletion of the FBM motif decreases the enrichment of BLTP2 at the tubular structures (D). (E) shows the method used for the quantification. The quantification result is shown in (F). Two-tailed t-test. Mean ±SEM. (FL) n=63 tubules from nine cells, (△FBM) n=111 tubules from 15 cells. (G) GFP-FBP17 localizes on the same BLTP2^Halo-positive tubules but does not overlap with the BLTP2^Halo signal. (H and I) GFP-mAmph2 co-localizes with BLTP2^Halo on the tubular structures (H), while GFP-mAmph2-BAR* (I) localizes on the same tubules but does not overlap with BLTP2 ^Halo. (J) AlphaFold3 predicts the interaction of a loop of BLTP2 (loop1) with the SH3 domain of amphiphysin. This loop harbors a RAPHPP sequence (RxP motif) which fits an SH3 domain binding consensus. (K and L) Co-expression of BLTP2^EGFP FL with a BLTP2^Halo construct in which each of these six amino acids of the SH3 binding consensus were mutated to alanine (BLTP2^Halo^RP→AA^), or BLTP2^Halo^RP→AA^ with the additional deletion of the FBM (BLTP2^Halo^RP→AA^-△FBM), showing a synergistic effect of abolishing SH3 domain binding and FAM102 binding in reducing the targeting of BLTP2 to the tubules. The quantification result is shown in (L). One way ANOVA. Mean ±SEM. (FL) n=63 tubules from nine cells, (RP→AA) n=103 tubules from 13 cells, (RP→AA and △FBM) n=97 tubules from 13 cells. (M) Schematic drawing depicting co localization of BLTP2, FAM102B and amphiphysin 2 on the PM-connected tubular endosomes, but segregation form FBP17.

### SH3 domain dependent interactions with endocytic BAR proteins cooperate with other factors in BLTP2 recruitment to PM-connected tubular endosomes

In our studies of HeLaM cells we noticed that although the enrichment of BLTP2^Halo-△FBM relative to full length BLTP2^EGFP at surface-exposed tubules was strongly reduced, it was not completely abolished (Figures 5D-5F). This implied that other factors may contribute to the enrichment of BLTP2 at the tubules. Tubular invaginations of the PM are sites where endocytic Bin-Amphiphysin-Rvs (BAR) domain containing proteins, for example FBP17 and amphiphysin family proteins, are known to assemble via the curvature sensing properties of their BAR domains^47–50^. Moreover, BAR domain containing proteins were implicated in the biogenesis of tubular recycling endosomes^51–54^ and were observed at Rab10-positive PM invaginations^27^. Typically, these proteins function as adaptors to recruit to curved bilayers, often via SH3 domains, a variety of other factors such as cytoskeletal scaffolds and signaling proteins^48,55,56^. Thus, we explored whether BAR proteins could contribute to the recruitment of BLTP2 at the distal portion of tubular recycling endosomes.

When we expressed the F-BAR domain proteins FBP17 (GFP-FBP17) and the N-BAR domain protein Amphiphysin 2 (GFP-mAmph2) (more precisely its non-neuronal isoform also referred to as BIN1) in HeLaM cells, both proteins, which contain C-terminal SH3 domains, became highly enriched on the BLTP2-positive tubular endosomes (Figures 5G and 5H). In agreement with previous studies of endocytic invaginations, GFP-FBP17, whose F-BAR domain has lower curvature than N-BAR domains, marked the portion of the tubules closer to the “outer” PM^57,58^ and was adjacent to, but did not overlap with, the BLTP2^Halo fluorescence (Figure 5G). In contrast, GFP-mAmph2, which associates with smaller diameter tubules^49,57^, was localized more deeply into the tubules and precisely colocalized with BLTP2 (Figure 5H). However, a construct of amphiphysin 2 lacking the SH3 domain (GFP-mAmph2-BAR*) still localized on the tubules but no longer colocalized with BLTP2 (Figure 5I), confirming an SH3 domain-mediated interaction between the two proteins, but also demonstrating that this interaction is not essential for the recruitment of BLTP2 to the tubules.

An interaction between BLTP2 and amphiphysin 2 was supported by overexpressing CFP-mAmph2 or GFP-Amph2-BAR* along with BLTP2^Halo in COS-7 cells that typically contain only few tubular recycling endosomes. Both overexpressed Amph2 and Amph2-BAR* induced PM tubular invaginations via their curvature generating properties, as expected^59^. While the tubules of this artificial system are very different in their origin and properties from tubular recycling endosomes, BLTP2 was recruited to them when they were generated by Amph2, but not by Amph2-BAR* (Figures S7A and S7B), supporting an SH3-mediated interaction. Consistent with the results in HeLaM cells, BLTP2^Halo was not recruited to tubules generated by GFP-FBP17 (Figure S7C).

In agreement with these results, AlphaFold3 predicted binding of the SH3 domain of mAmph2 to an amino acid loop of BLTP2 that projects out of its rod-like core near its N-terminus (pTM=0.55) (Figure 5J). This loop contains the core SH3 binding consensus “RxP” motif^60^ within the “RAPHPP” sequence which is conserved among mammalian BLTP2s (Figure S7D). When a BLTP2 construct in which all these 6 amino acids had been mutated to alanine (BLTP2^Halo^RP→AA^) was expressed in HeLaM cells, its enrichment on the tubules relative to the enrichment of BLTP2^EGFP-FL was reduced (Figures 5K and 5L). Moreover, combining this mutation with the deletion of the FBM (BLTP2^Halo^RP→AA^-△FBM) almost completely abolished its enrichment on the tubules (Figures 5K and 5L). We conclude that both FAM102B and SH3-domain N-BAR domain proteins contribute to recruit BLTP2 to PM-connected tubular endosomes (Figure 5M).

### BLTP2 is recruited to macropinosomes undergoing fusion with PM

In the course of our live imaging experiments involving cell expressing fluorescently tagged BLTP2, we sometimes observed transient flashes of focal BLTP fluorescence close to the cell surface. When these experiments were performed in the presence of anti-MHC-I antibodies to label broadly the PM and macropinosomes, we found that these flashes correspond to the exocytosis of macropinosomes. This was clearly exemplified by experiments in COS-7 cells, which have only few focal BLTP2^Halo accumulations at ER-PM contacts (unless co-expressed with FAM102) and therefore allows easy visualization of these events.

Most macropinosomes, shortly after internalization, acquired as expected markers of early endosomal stations, such as Rab5, its effector APPL1 and PI3P^61–63^. However, a subset of them did not progress to this stage and instead acquired bright transient spots of BLTP2 signals, revealing formation at their surface of contacts with the ER where BLTP2 is highly concentrated. Strikingly, the appearance of BLTP2 on these macropinosomes correlated with a major change of their shape. They shrank and transformed from vacuolar structures into tubules that appeared to be connected to the PM while remaining BLTP2 positive. Eventually these structures disappeared, suggesting their collapse into the PM, with a corresponding loss of the BLTP2 signal (Figures 6A, S8A and S8B; Videos S6 and S7).

**Figure 6.**
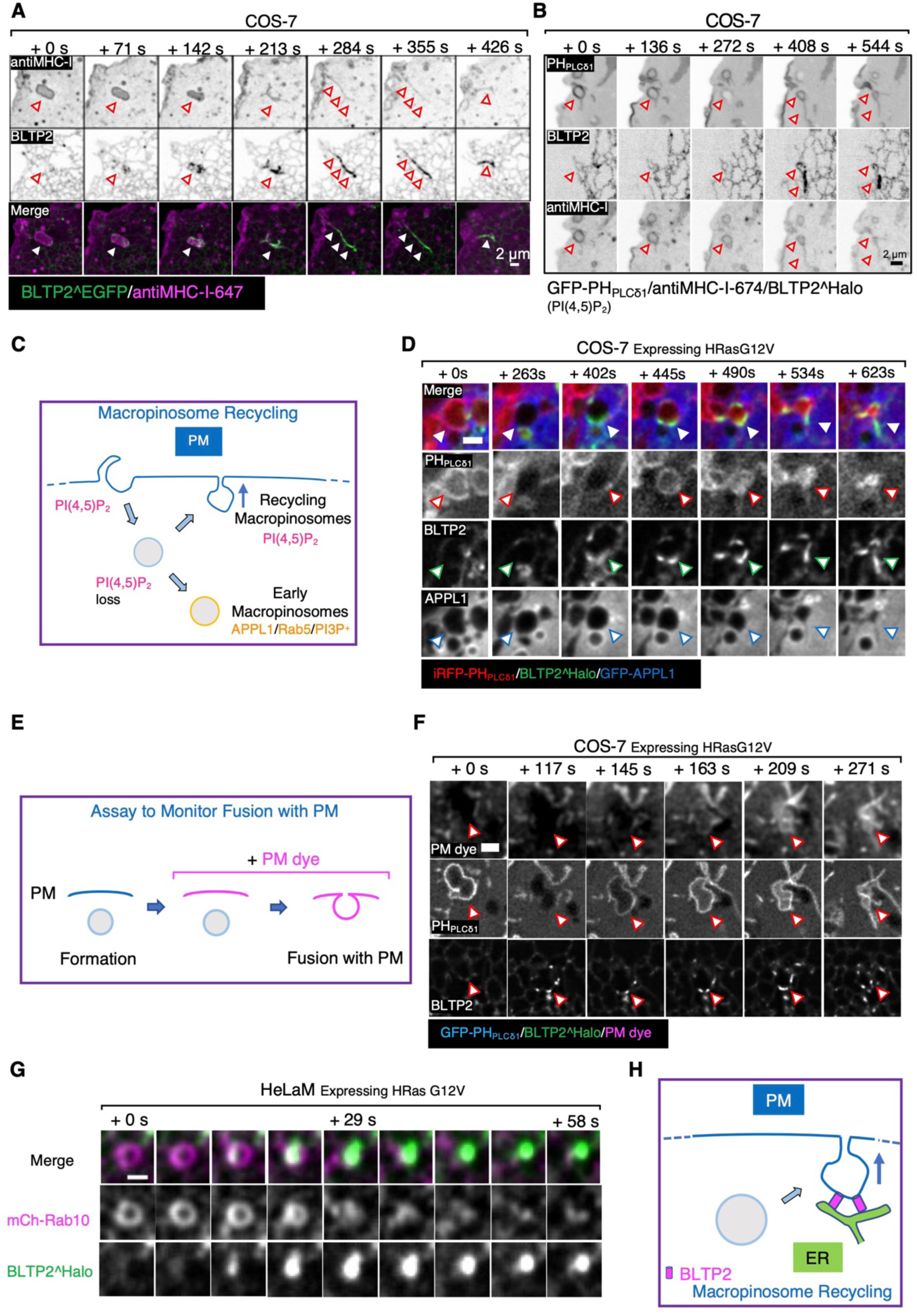
BLTP2 is recruited to recycling macropinosomes undergoing fusion with the PM. (A and B) COS-7 cells showing macropinosomes (labeled by internalized fluorescent anti MHC-I antibodies) that undergo a dramatic morphological change and acquire BLTP2^EGFP as they fuse and collapse with the PM. In (B), a newly formed macropinosome at first loses PI(4,5)P_2_, but then reacquires PI(4,5)P_2_ as it fuses with the PM. (C) Schematic drawing of macropinosome recycling. A nascent macropinosome first loses PI(4,5)P_2_ after fission from the PM, then it acquires the identity of early endosomes (PI3P, APPL and Rab5), or regain PI(4,5)P_2_ as it fuses back to the PM. (D) Stimulation of macropinocytosis by expression of HRas G12V in COS-7 cells. BLTP2^Halo is recruited to a newly formed macropinosome that loses PI(4,5)P_2_ but does not acquire APPL2 and regains PI(4,5)P_2_ signaling during its recycling back to the PM. (E) Schematic drawing depicting the assay to monitor macropinosome fusion with the PM. PM dye is added after the formation of macropinosomes. Pre-formed macropinosomes will not be labeled by the dye until they fuse with the PM. (F) A BLTP2-positive macropinosome gains access to the PM dye showing that it fuses with the PM. Macropinocytosis was stimulated by expressing HRas G12V. (G) Acute recruitment of BLTP2^Halo to a macropinosome in the process of fusing with the PM in HeLaM cells. The macropinosome is also positive for mCherry-Rab10. Scale Bar 1μm. (H) Schematic drawing of the recruitment of BLTP2 to a recycling macropinosome undergoing fusion with the PM.

The fusion of these organelles with the PM was confirmed by monitoring the dynamics of PI(4,5)P_2_, a defining lipid of the PM^29^, on their surface. In cells expressing the PI(4,5)P_2_ probe PH_PLCδ1_, PI(4,5)P_2_ disappeared from nascent macropinosomes as expected, as PI(4,5)P_2_ is known to be rapidly removed from endocytic vesicles, primarily via the action of PI(4,5)P_2_ phosphatases^64^. However, on the macropinosomes which had acquired BLTP2 signal, a concomitant resurgence of PI(4,5)P_2_ was observed (Figure 6B; Video S8), as expected if they had fused with the PM and thus could acquire this phospholipid via its diffusion from the surrounding PM bilayer. Most likely, acquisition of PI(4,5)P_2_ is a key signal that triggers formation of BLTP2-dependent ER-PM tethers on the macropinosomes that undergo exocytosis.

Similar results were obtained in COS-7 and HeLaM cells where macropinocytosis was induced by the expression of the constitutively active mutant form of HRas (HRas G12V)^65^. Even in these cells we found that after losing PI(4,5)P_2_, some macropinosomes, which were labeled by the PI(3,4,5)P_3_/PI(3,4)P_2_ marker PH_AKT_ as expected^66–68^, did not mature into the APPL1 or Rab5 stage (Figures 6D, S9A and S9B). Instead they retained the PI(3,4,5)P_3_/PI(3,4)P_2_ signal and then showed BLTP2 recruitment correlated with shrinking (Figures S9A). To confirm that PI(4,5)P_2_ resurgence reflected fusion events with the PM, we added to cells the non-permeable membrane dye CellBrite® steady 650 that cannot have access to macropinosomes generated before its addition (Figure 6E). Observation of these cells showed that resurgence of PI(4,5)P_2_ on macropinosomes coincided with their labeling by CellBrite® steady 650, thus indicating opening of their lumen to the cell surface (Figure 6F; Video S9). Finally, macropinosomes that became positive for PI(4,5)P_2_ and BLTP2 were also positive for Rab8 and Rab10, two RabGTPases implicated in exocytosis of a variety of vesicles^69–72^, including macropinosomes^73^ (Figures 6G and S9C).

The occurrence of macropinosomes that fail to acquire early endocytic markers, acquire instead Rab8 and Rab10, and recycle back to the PM is in agreement with other studies^73,74^. It was also reported that the proportion of these recycling events is strongly increased upon inhibition of VPS34^73^, the kinase that generates on early endosomes PI3P, the signature phosphoinositide of these organelles. Accordingly, upon treatment of COS-7 cells with the VPS34 inhibitor SAR405^73^ we observed many PI3P positive macropinosomes that re-acquired PI(4,5)P_2_, became positive for BLTP2 and fused with the PM (Figure S9D).

### Absence of BLTP2 impairs the collapse of macropinosomes into the PM after their fusion

Collectively, the results reported above suggest that a primary localization of BLTP2 is at contacts between the ER and the PM. Such localization, in turn, suggests that BLTP2-dependent transport of lipids from the ER to the PM may be required to define and maintain the properties of the PM, with an important impact on its dynamics. To address this possibility, we generated BLTP2-KO HeLaM cells using CRISPR-Cas9 based strategy. Single KO clones were isolated and verified both by nucleotide sequencing of the edited region and by Western blotting (Figure 7A).

**Figure 7.**
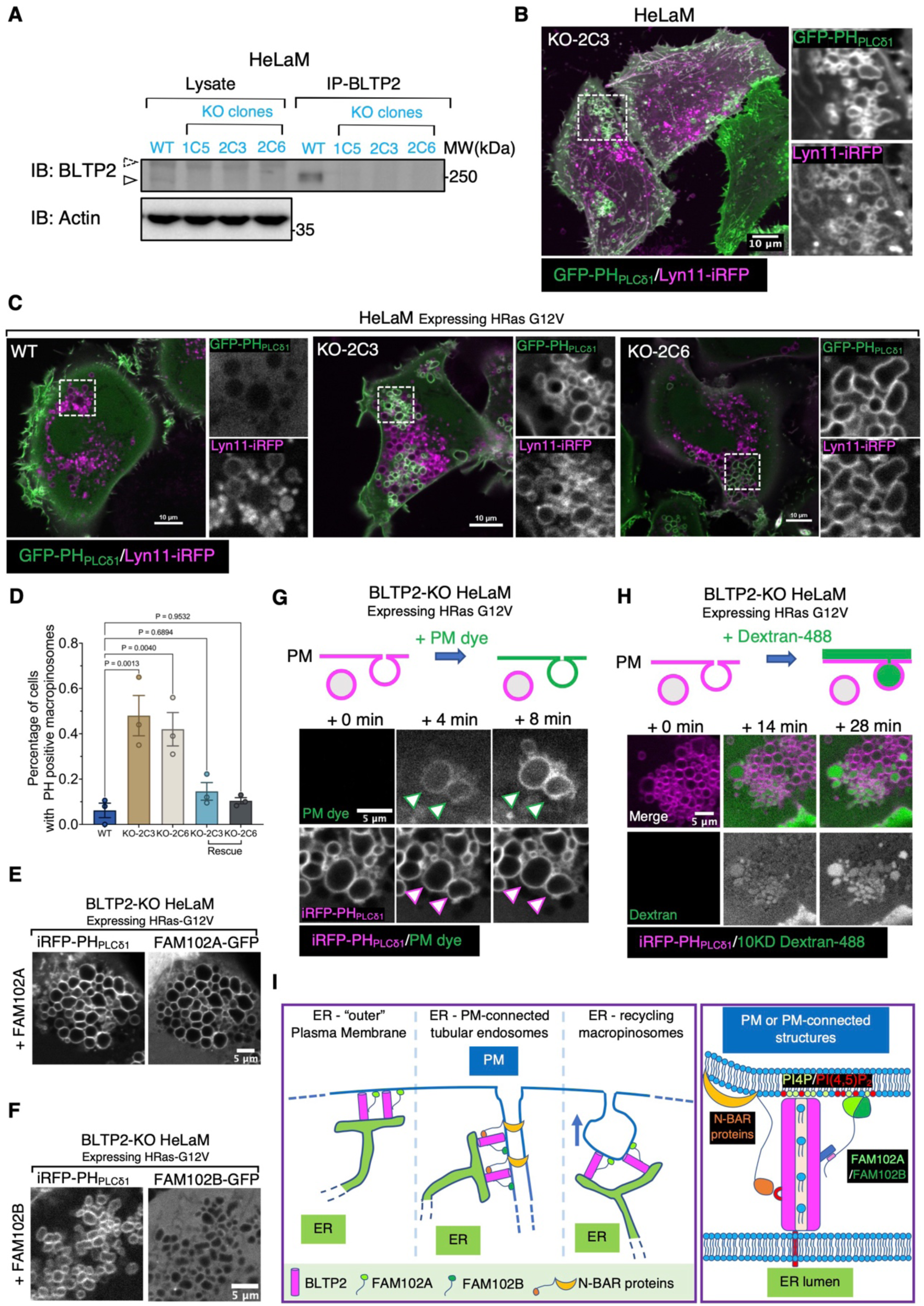
BLTP2-KO cells show accumulation of intracellular PI(4,5)P_2_-positive vacuoles. (A) Western blot validating knock-out (KO) of BLTP2 in HeLaM cells. Endogenouse BLTP2 is enriched using IP before detection. Three independent KO clones are verified. (B) BLTP2-KO HeLaM cells showing presence of PI(4,5)P_2_-positive intracellular vacuoles. Tubular recycling endosomes are still present in these cells. (C and D) Expression of HRas G12V in HeLaM cells (two independent clones: KO-2C3 and KO-2C6) induces formation of macropinosomes/intracellular vacuoles in both WT and BLTP2-KO cells. However, only in the KO cells a large fraction of these vesicles remain PI(4,5)P_2_ positive. Quantification of PI(4,5)P_2_ positive macropinosomes is shown in (D). One way ANOVA. Mean ±SEM. n=3 independent experiments. 123 cells for WT, 137 cells for KO-2C3, 161 cells for KO-2C6,119 cells for KO-2C3 rescue and 122 cells for KO-2C6 rescue. (E and F) BLTP2-KO cells expressing HRas G12V together with FAM102A-GFP (E), or FAM102B-GFP (F). FAM102A-GFP, but not FAM102B-GFP, is enriched on PH_PLCδ1_ labeled PI(4,5)P_2_ positive macropinosomes. (G and H) PM dye (G), or 10KD Dextran-488 (H) were added to BLTP2-KO cells expressing HRas G12V. A fraction of the pre-existing PI(4,5)P_2_ positive vacuoles were labeled by the dye within minutes after PM dye addition (G), or showing internalization of Dextran-488 (H). (I) Schematic model of BLTP2 localization at contacts of the ER with PM and PM-connected structures (left). Illustration of the molecular interaction of BLTP2 with phosphoinositides and its binding proteins (FAM102A/B, N-BAR domain proteins) at these contact sites (right).

Inspection of WT and BLTP2 KO cells revealed formation of PI(4,5)P_2_-positive (as revealed by PH_PLCδ1_ labeling) vacuoles in the KO cells, while the PM-connected tubular endosomes seem undisrupted (Figure 7B). However, when expressing constitutively active RAS (HRas G12V), not only most KO cells showed a massive accumulation of macropinosomes relative to controls, but many of these macropinosomes, in contrast to those present in WT cells, were positive for PI(4,5)P_2_ (Figures 7C and 7D) and for FAM102A (Figure 7E), but not FAM102B (Figure 7F). Additionally, many of these PI(4,5)P_2_ and FAM102A positive vacuoles were connected to the PM, as they were accessible to both the non-permeable membrane dye CellBrite® steady 650 (Figure 7G; Video S10) and to 10kD dextran-488 (Figure 7H; Video S11). Importantly, this phenotype was rescued by exogenous expression of BLTP2 (Figure 7D), confirming its BLTP2-dependence. Based on these observations, we suggest that these vacuoles represent post-fusion structures whose collapse into the PM is impaired, or macropinosomes that failed to undergo fission from the PM. Irrespective of the mechanisms underlying their formation, these results indicate that loss of BLTP2 has a major impact on the dynamics of the PM.

## Discussion

Our study shows that BLTP2 is primarily concentrated at contacts between the ER and the PM in commonly used mammalian cell lines, although with cell-specific features, and identifies some of the interactions responsible for these localizations. We also show that the absence of BLTP2 results in the presence of intracellular vacuoles with PM-like properties and in least in some cases still connected to the PM, consistent with a role of this protein in controlling the dynamics of the cell surface, most likely via its properties to transport phospholipids from the ER to the PM.

Our findings reconcile discrepancies from the previous literature which had reported a localization of BLTP2 orthologues at ER-PM contacts in yeast, drosophila and mammalian cell lines^12,17^, but a selective localization at contacts between the ER and tubular recycling endosomes in another study of mammalian cells^18^. We confirmed that the tubules reported by this study have the reported properties of tubular recycling endosomes, as they are positive for Rab8 and Rab10 as well as for proteins, such as MHC-I, found to populate such endosomes. However, we found that, when positive for BLTP2, these tubular membranes are connected to the PM and thus represent another example of ER-PM localization. The connection to the PM of Rab10-positive tubules were also recently reported in a study which describes these tubules as PM invagination^27^. These tubules are very abundant in HeLaM cells, where they are typically very long, often extending from the Golgi complex region to the PM. At least in most cases these tubular structures do not appear to be typical transport intermediates, but lasting structures. Moreover, they appear to be heterogeneous in properties along their length, as they have *bona fide* PM properties selectively on their portions closer to the outer cell surface, as exemplified by the formation of BLTP2-dependent tethers and by the presence of other classical ER-PM tethers, such as TMEM24 and MAPPER, selectively in these regions. Similar tubules with an enrichment of contacts with the ER at their surface were described by Yao et al^75^. It is possible that a special concentration of BLTP2 and other lipid transport proteins at these sites may help ensuring PM-like lipid composition on the distal portions of the tubules which are directly adjacent to, and continuous with, the “outer” plasma membrane.

Other sites where we have detected presence of BLTP2-positive contacts are macropinosomes in the process of fusing with the PM. These are macropinosomes that fail to migrate and mature to late stations in the endocytic pathway and instead recycle back to the PM. As they fuse with the PM, such macropinosomes undergo a dramatic shrinkage and remodeling of their membrane which correlates with the recruitment of BLTP2 positive contacts. BLTP2-mediated lipid transport at this stage may help provide lipids for this remodeling or to make the membrane of macropinosome compatible with its intermixing with the PM. The appearance of flashes of BLTP2 fluorescence on macropinosomes (reflecting acute formation of BLTP2-positive ER-PM contacts) only when they have established continuity with the PM, provides a striking demonstration of the PM binding specificity of BLTP2.

Concerning mechanisms through which ER-anchored BLTP2 binds *in trans* the PM to achieve bridge-like lipid transport, we have found evidence for both lipid-based and protein-based interactions. We have shown that phosphoinositides in the PM are important. Moreover, we have identified FAM102A and FAM102B as PM-associated adaptor proteins that bind BLTP2 via an interaction conserved from yeast to mammals, a discovery independently made by Elizabeth Conibear and co-workers who found that the yeast homolog of FAM102A/B (Ybl086c) binds to, and targets, yeast BLTP2 to ER-PM contacts (personal communication). Clearly, the overexpression of these two proteins enhances BLTP2 targeting to the PM. Conversely, removal of the FAM102 binding motif from BLTP2 strongly decreases, although does not abolish, its PM targeting, indicating that FAM102 proteins are not the only determinants of BLTP2 localization. While these two proteins, which comprise an N-terminal C2 domain followed by an approximately 240 a.a. long predicted unfolded region, are very similar to each other, they also have some different properties, as only FAM102B selectively accumulates with BLTP2 at the tubular invaginations of the PM which are continuous with tubular recycling endosomes. Interestingly, the yeast interactome database which reports a high confidence interaction of the yeast BLTP2 ortholog Fmp27 with the yeast Fam102 protein (Ybl086c), also identifies Osh3, Ist2 and Lro1 as components of the Fmp27-Ybl086c network (Figure 4A). Both Ist2 (TMEM16 in mammals) and Osh3 (a member of the mammalian ORP family) are proteins implicated in lipid dynamics at ER-PM contacts^40,41^, while Lro1 (LCAT in mammals), is a phospholipid metabolizing enzyme^42^, raising the possibility of a functional cooperation with these proteins with BLTP2. Finally, we found evidence for SH3 domain-dependent interactions of BLTP2 with endocytic BAR domain proteins, such as amphiphysin 2, which may help better explain the concentration of BLTP2 on the tubules.

A recent study reported that BLTP2 and its orthologues in yeast play a role in controlling the appropriate fluidity of the PM bilayer by preferentially delivering phosphatidylethanolamine from ER^12^. How BLTP2 could control a specific enrichment of this phospholipid in the PM remains unclear, but an impact of BLTP2 on PM fluidity could be one explanation for the abundant presence of intracellular vacuoles connected to the PM in BLTP2-KO cells, more so in cells expressing HRas G12V to stimulate bulk endocytosis. This accumulation may reflect impairment of the ability of this membrane to appropriately remodel in response to incoming and outgoing membrane traffic. Ectopic PI(4,5)P_2_ on intracellular membrane was also observed in Drosophila BLTP2 mutant cells^17^, although the potential connection of these vacuoles to the PM was not explored. Another BLTP, BLTP1, which shares special similarities to BLTP2^4,7^, including the presence of a transmembrane region anchored to the ER, was also reported to control fluidity of PM, in this case by ensuring delivery of phospholipid with appropriate fatty acid composition (primarily saturation level of their aliphatic chains)^7,13–15^. The importance of both BLTP1 and BLTP2 for organismal life, proven by early embryonic lethality of KO mice and by the developmental defects of BLTP1 and BLTP2 Drosophila mutants^9,76^ are striking demonstrations that direct protein-mediated bulk lipid transport from the ER to the PM, a process unknown until recently, has a general and fundamental importance in cell physiology.

## Resource availability

### Lead contact

Further information and requests for resources and reagents should be directed to the lead contact, Pietro De Camilli.

### Materials availability

Plasmids and all other reagents generated in this study are available upon request from the lead contact.

### Data and code availability

This paper does not report original code. Any additional information is available from the lead contact upon request.

## Acknowledgements

We thank Elizabeth Conibear for sharing information about the interaction of BLTP2 with FAM102 family proteins before publication. We also thank Benjamin Johnson, Hongyan Hao and Hely Rodriguez Cruz (De Camilli Lab) for discussion, Chuan Liu and Anthony O’Donnell (Johns Hopkins University) for help and discussion with structural predictions, Alina Vulpe for technical support. This work was supported by the NIH grants and fellowships: DA018343 and NS036251 to P.D.C., F31NS135930 and T32GM145469 to C.A., T35DK104689 to J.N.E and by a Gruber Science Fellowship to H.Y.

## Author contributions

A.D. and P.D.C. conceptualized the project. K.F. and Y.W. performed the correlative light electron microscopy experiments. A.D. performed all other experiments reported in this study with contributions from P.X., C.A. and H.Y.. C.A., J.N.E. and A.G-S. made the initial observation leading to this study. A.D. wrote the original draft of the manuscript with the input from all other authors. P.D.C. supervised the project and wrote the manuscript along with A.D.

## STAR Methods

### Key resources table

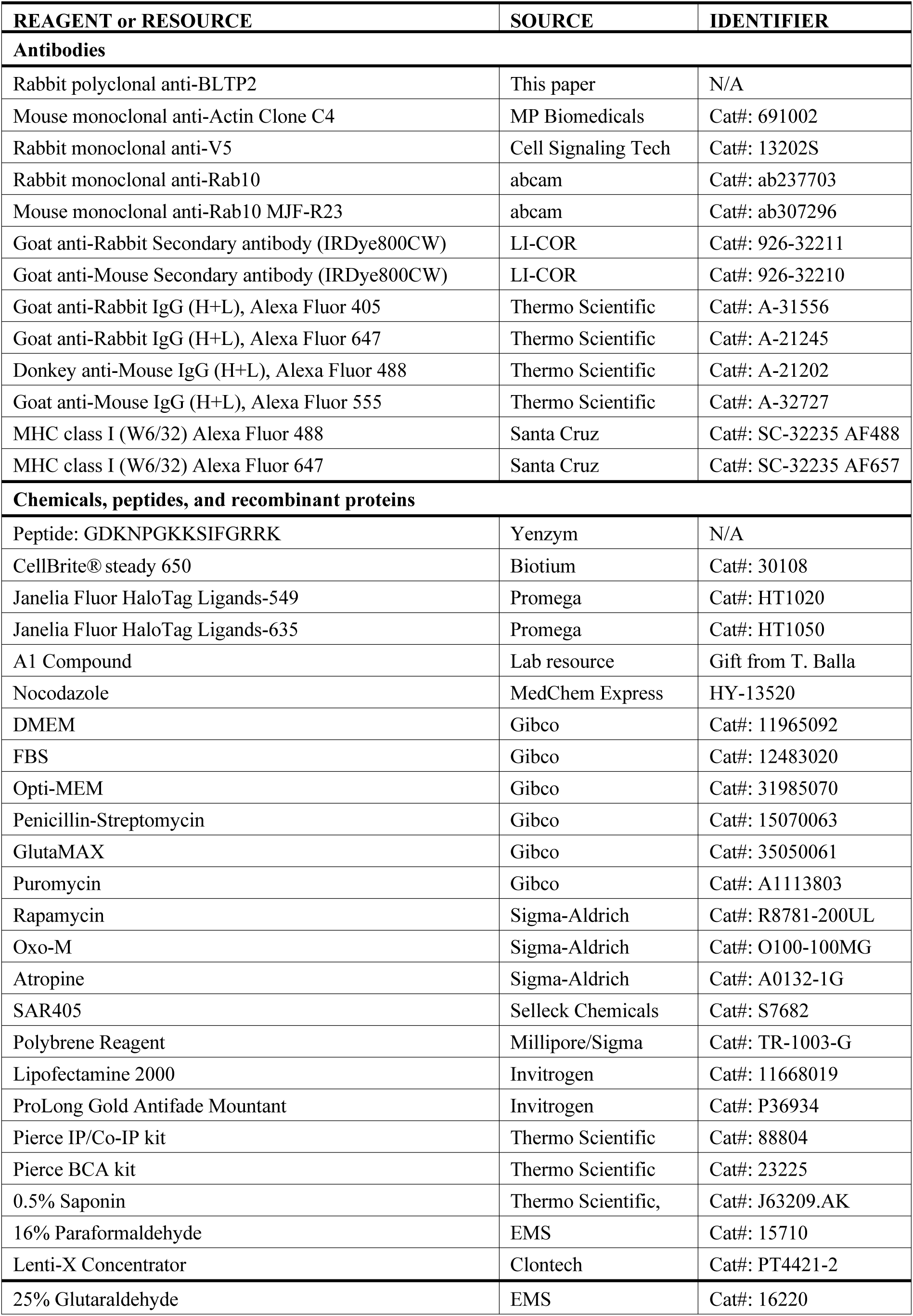
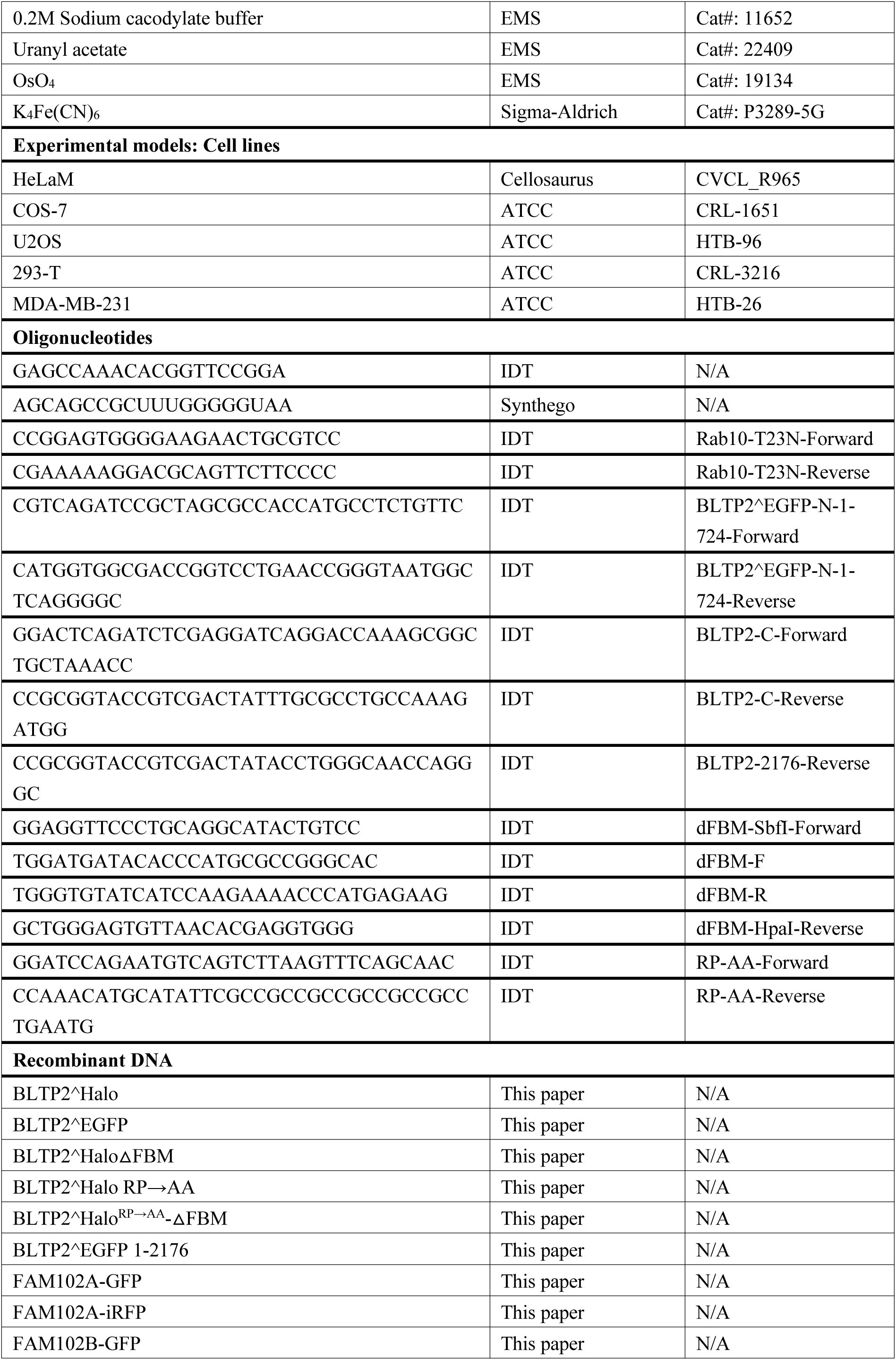
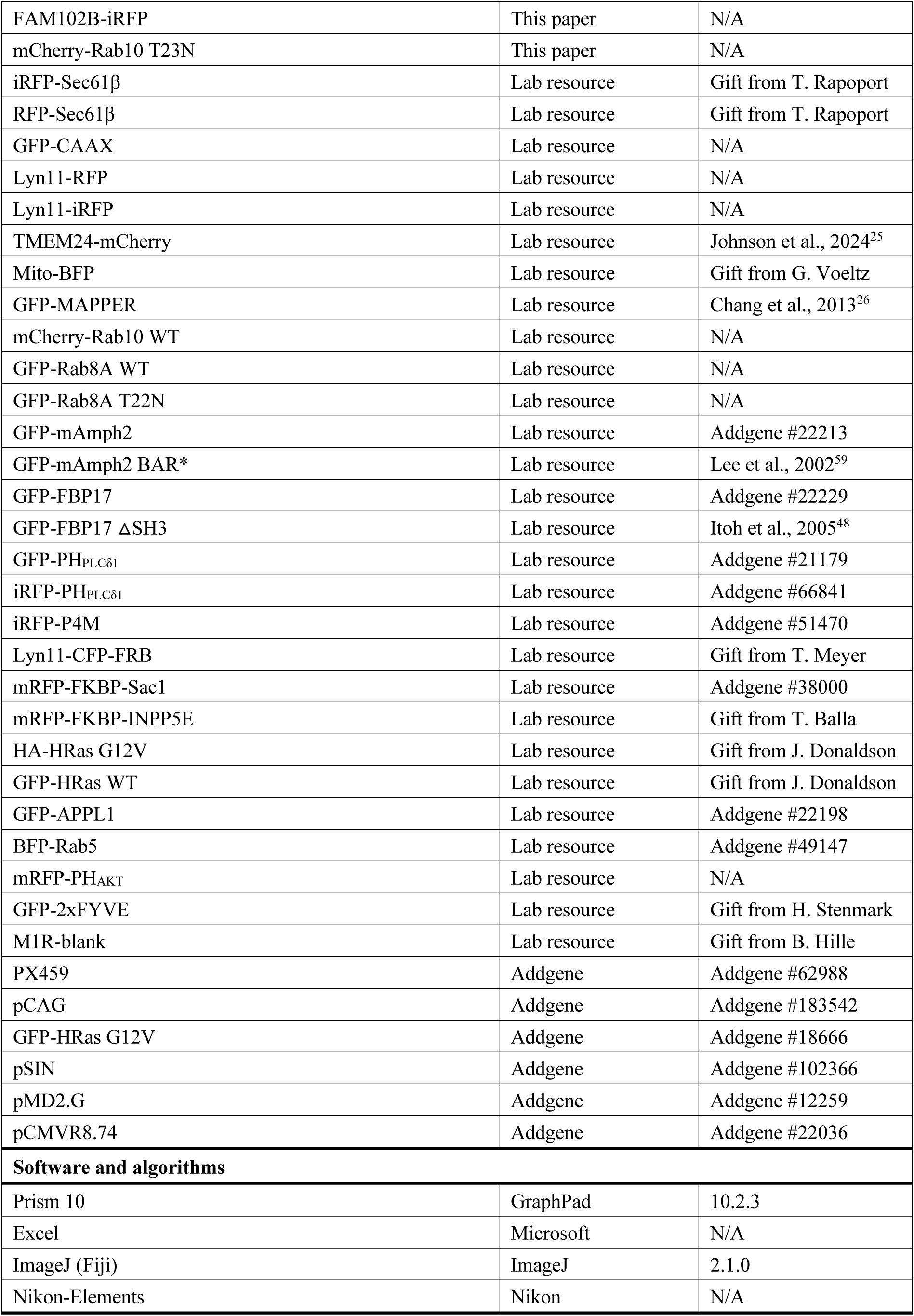

### Plasmids

The original clone containing the BLTP2 ORF (NCBI reference sequence: NM_014680.5) was obtained from GenScript. Internally tagged BLTP2^EGFP was generated by first amplifying the N-terminal (a.a.1-724) and C-terminal (a.a.725-2235) fragments of BLTP2 using PCR, respectively. The N-terminal fragment was inserted into a pEGFP-C1 plasmid between the NheI and AgeI restriction sites using In-Fusion system (Takara) to create an intermediate construct “Nterm-1-724_EGFP”. The C-terminal fragment was then inserted in this intermediate construct between the XhoI and SalI restriction sites to create the full BLTP2^EGFP construct. For BLTP2^Halo, EGFP was replaced with a Halo tag. The full BLTP2^EGFP or BLTP2^Halo sequence was further amplified and inserted between the XbaI and EcoRI restriction sites in a pCAG vector using In-Fusion system (Takara), respectively.

BLTP2 C-terminal truncation constructs were generated by amplifying each fragment and inserting them into the “Nterm-1-724_EGFP” construct. BLTP2^Halo-△FBM was generated using over-lap PCR to remove the FBM sequence. The BLTP2^Halo^RP→AA^ and BLTP2^Halo^RP→AA^-△FBM mutation constructs were generated using over-lap PCR to replace the region to be mutated.

Both FAM102A (NCBI reference sequence: NM_001035254.3) and FAM102B (NCBI reference sequence: NM_001010883.3) cloned in a pcDNA3.1-C-eGFP vector between the KpnI and BamHI restriction sites were obtained from GenScript. FAM102A-iRFP and FAM102B-iRFP were generated by swapping the EGFP with iRFP between the NotI and BsrGI restriction sites. mCherry-Rab10 T23N was generated using over-lap PCR to introduce the point mutation. All primers and other constructs used in this study were listed in the Key resource table.

### Cell culture and transfection

All cell lines were cultured at 37℃ and 5% CO_2_ in DMEM supplemented with 10% Fetal Bovine Serum (FBS), 1x Penicillin-Streptomycin and 1x GlutaMAX. For transient transfection, cells were first seeded in a 35 mm glass bottom dish (MatTek). When cells reached 60-80% confluency, 1-3 μg plasmids and 2 μl Lipofectamine 2000 were diluted in pre-warmed Opti-MEM for 5 min, respectively, and then mixed for another 15 min before their addition to the cells.

### Generation of BLTP2 stable cell lines using Lentivirus

BLTP2^Halo or BLTP2^EGFP were cloned into a pSIN vector, respectively. This vector was mixed with packaging vectors pMD2.G and pCMVR 8.74 and transfected into 293-T cells for virus production. 24 hours post transfection, fresh medium was added to the cells. Medium containing the viruses was collected after another 24 hours. Viruses were concentrated using Lenti-X concentrator following manufacture’s protocol by mixing 1x volume of the concentrator with 3x volume of clarified medium (by passing through 0.45 μm filter) and incubate at 4 ℃ overnight. Pellets were then collected after centrifugation and resuspended using PBS. Virus suspension was either used directly or further stored in -80 ℃. For viral transduction, viruses were mixed with Polybrene reagents and added to the cells. 48 hours after adding the viruses, cells were incubated in the presence of 2 μg/ml puromycin for 5 days for selection. Single clones were isolated by serial dilution and cultured in 96-well plates. Positive clones were verified by fluorescence and expanded.

### CRISPR-Cas9 based generation of BLTP2 knock-out and knock-in cell line

For BLTP2-KO cell line, PX459 vector containing a guide RNA sequence was transfected into target cells (HeLaM). 24 hours post transfection, cells were treated with 1 μg/ml puromycin for 5 days. Single cell was isolated using serial dilution and cultured in 96-well plates, then expanded in 24-well plates. Genomic DNA was extracted, and successful editing was first verified using PCR and sequencing, then further verified using western blot. The BLTP2^2xV5-KI cell line was generated by Synthego. In brief, a ribonucleoprotein with guide RNA and spCas9 was delivered with a donor sequence to the cells (HeLaM). Single clones were isolated and validated using PCR followed with sequencing. Final clones used in this study were further validated using western blot.

### Immunoprecipitation

Immunoprecipitation of BLTP2 was performed using the Pierce IP/Co-IP kit following the procedures indicated by the manufacturer. Briefly, cells were first washed with PBS and then directly lysed by adding ice-cold IP lysis buffer to the culture dish (500 μl buffer per one 10-cm dish). Lysed cells were scraped and transferred into a 1.7 ml tube. Cells were further lysed under rotation at 4 ℃ for 30 min. Cell lysate was clarified using a tabletop centrifuge at 17,000 g for 30 min. 8 μg of BLTP2 antibody was added to the supernatant and incubated overnight while rotating at 4 ℃. The immuno-complex was then enriched using protein A-G coated magnetic beads by incubation at room temperature for 2 hours. The beads were washed 2X with IP lysis buffer followed with one wash in H_2_O. Bound proteins were then eluted with “elution buffer” (pH 2.0) and neutralized with “neutralization buffer” (pH 8.5). After mixing with SDS loading buffer, samples were directly processed for SDS-PAGE without heating.

### Microscopy

All imaging experiments were performed using either a CSU-W1 Sora (Nikon) or a Dragonfly spinning-disk confocal imaging system (Andor). For live imaging, cells were maintained in DMEM at 37℃ and 5% CO_2_ during imaging. In cells involving Halo constructs, Janelia Fluor HaloTag ligands were added to the culture medium at 1:4000 dilution for 10 min. This medium was then replaced by fresh medium not containing Halo ligand before imaging. For immunofluorescence microscopy, cells were fixed using 4% paraformaldehyde (PFA) for 10 min at room temperature. Fixed cells were then permeabilized and blocked using 0.05% Saponin in 10% FBS for 1 h and incubated with primary antibody at 4 ℃ overnight, followed by secondary antibody at room temperature for 1 h. Finally, cells were mounted using ProLong Gold Antifade Mountant and kept at 4 ℃ before imaging.

To stimulate macropinocytosis cells were transfected with HRas G12V and imaged 48 hours post transfection. In some experiments the medium was replaced with fresh medium containing PM dye (CellBrite) (1:4,000) during imaging. <u>F</u>luorescent labeled MHC-I antibodies were diluted in cell culture medium (final concentration 20 μg/ml) which was then added to the cells. Cells were imaging live by confocal microscopy within 30 min.

#### Manipulations to deplete PI4P or PI(4,5)P_2_ at the PM

To reduce PI4P in the PM, the PI4KIIIα inhibitor A1 was diluted into prewarmed culture medium to a concentration of 200 nM, then added to the cells at 1:1 ratio at the start of the imaging session (final concentration 100 nM). For the recovery, one hour after A1 treatment culture medium was removed and cells were washed twice in PBS, then changed into fresh medium for further imaging. To acutely deplete PI(4,5)P_2_, cells were transfected with a plasmid encoding M1R with no fluorescent tag (M1R-blank). The M1R ligand Oxo-M diluted in culture medium to a concentration of 20 μM was added to cells after the start of live imaging at 1:1 ratio (final concentration at 10 μM). Eight mins after addition of Oxo-M, the M1R antagonist atropine (final concentration at 50 μM) was added to reverse the stimulation.

### Correlative Light and Electron Microscopy (CLEM)

For CLEM, HeLaM cells were plated on 35 mm MatTek dish (P35G-1.5-14-CGRD) and transfected as described above with plasmids encoding BLTP2^Halo and Mito-BFP. Cells were fixed with 4% PFA in PBS, then washed three times with PBS before being analyzed by fluorescence light microscopy imaging. Regions of interest were selected and their coordinates on the dish were identified using phase contrast. Cells were further fixed with 2.5% glutaraldehyde in 0.1 M sodium cacodylate buffer, postfixed in 2% OsO_4_ and 1.5% K_4_Fe(CN)_6_ in 0.1 M sodium cacodylate buffer, *en bloc* stained with 2% aqueous uranyl acetate, dehydrated, and embedded in Embed 812. Cells of interest were relocated based on the pre-recorded coordinates. Ultrathin sections (50-60 nm) were observed in a Talos L 120C TEM microscope at 80 kV, images were taken with Velox software and a 4k × 4K Ceta CMOS Camera (Thermo Fisher Scientific).

### Image processing and statistical analysis

Image processing was performed using either ImageJ (Fiji) or the Nikon Elements software package with built-in plugins. Statistical analysis was performed using Prism 10 (GraphPad).

---

**Figure S1.**
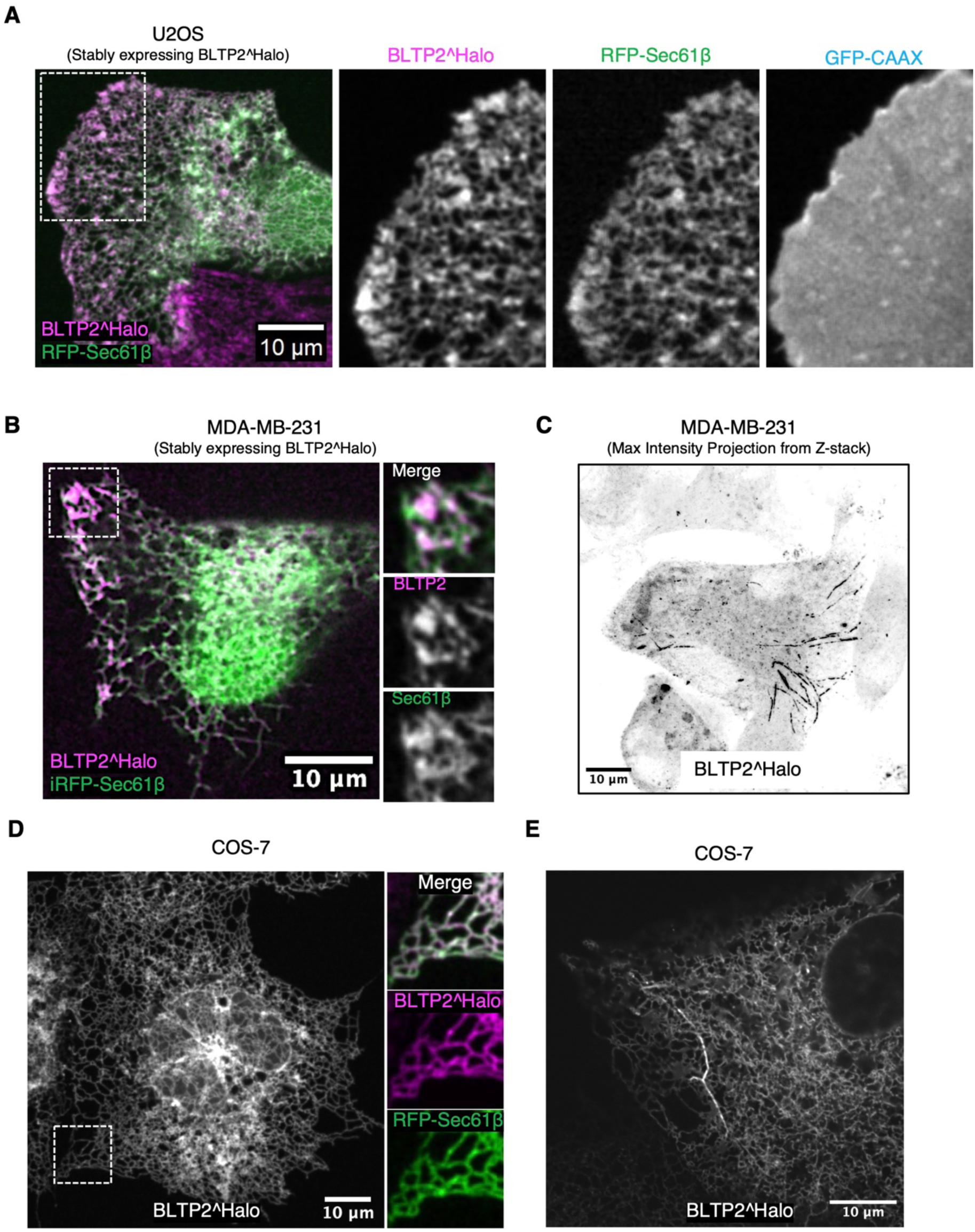
BLTP2^Halo show heterogeneous localization in different cell types. (A and B) U2OS (A) or MDA-MB-231 (B) cells stably expressing BLTP2^Halo show its localization throughout the ER (labeled with ER marker Sec61β) with an enrichment at ER-PM contacts. (C) In some MDA-MB-231 cells, BLTP2^Halo show an enrichment on tubular structures similar to those observed in HeLaM cells. (D and E) BLTP2^Halo mainly localizes throughout the ER in COS-7, as shown by colocalization with the ER marker RFP-Sec61β (D), and only occasionally it is enriched at some tubular structures (E).

**Figure S2.**
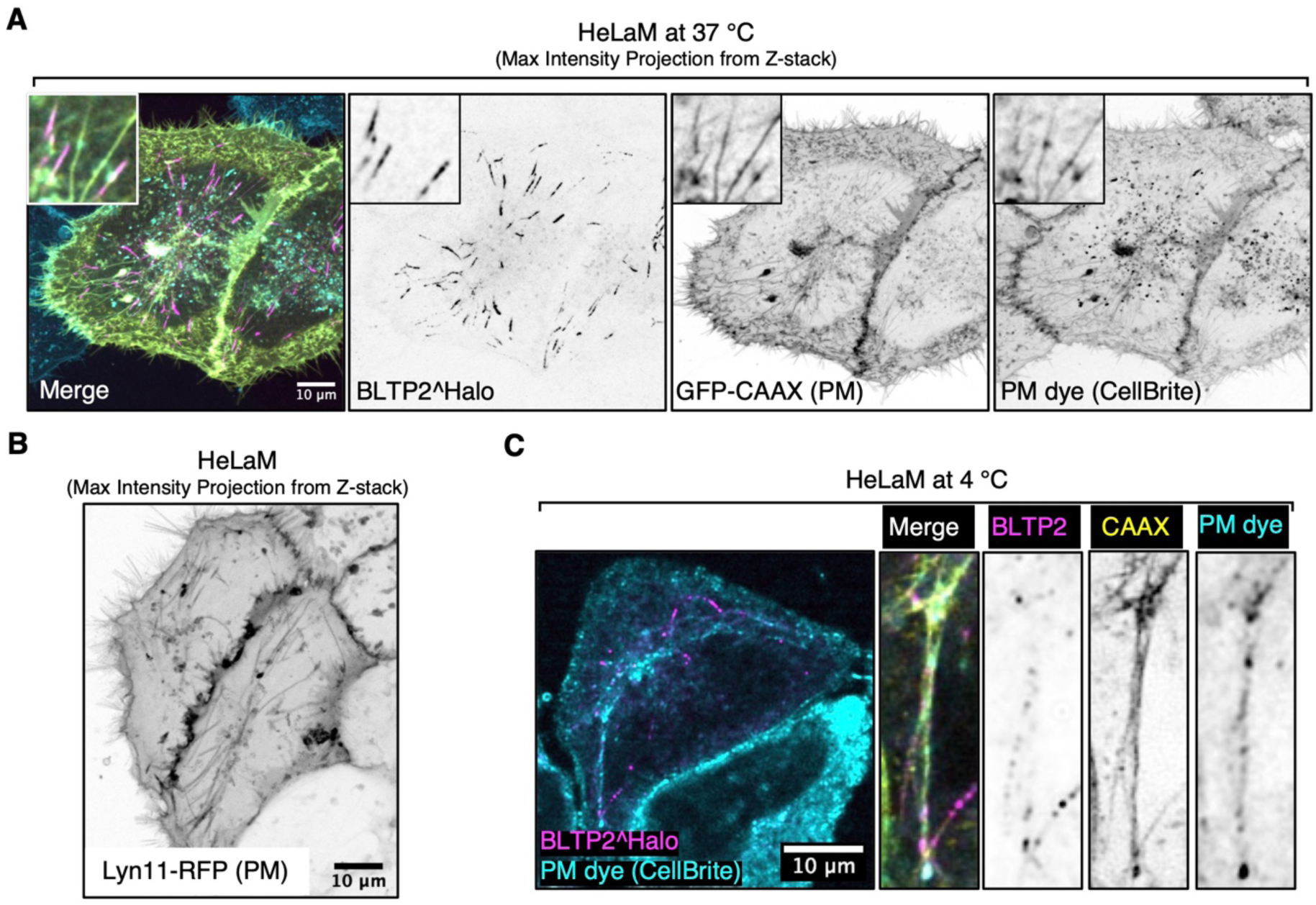
BLTP2-positive tubular endosomes are positive for PM markers. (A) BLTP2^Halo is enriched at tubular structures positive for the PM marker GFP-CAAX and for an extracellular membrane impermeable dye CellBrite labeled at 37 ℃. (B) The tubular structures of HeLaM cells are also positive for another PM marker, Lyn11-RFP. (C) BLTP2-positive tubules are also labeled by CellBrite at 4 ℃.

**Figure S3.**
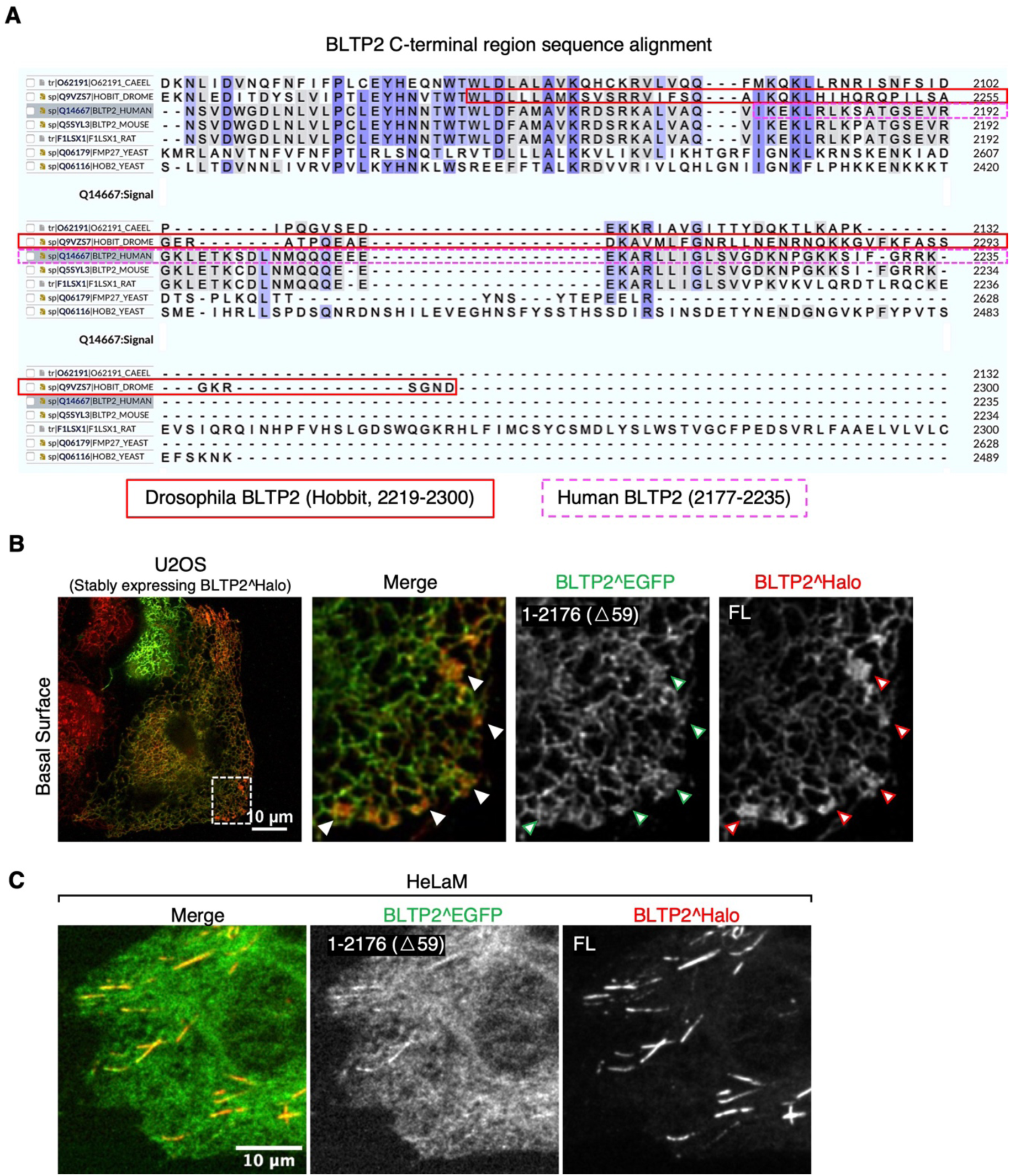
The C-terminal region of BLTP2 is required for its tethering function. (A) Sequence alignment of the C-terminal region of human BLTP2 with the corresponding region of other species. Conserved residues are highlighted in blue, with darker color indicating higher conservation. a.a.2219-2300 of Drosophila BLTP2 (hobbit) are framed by red solid lines and a.a.2177-2235 of human BLTP2 by magenta dashed lines. (B and C) U2OS cells (B) and HeLaM cells (C) co-expressing the BLTP2 C-terminal deletion mutant BLTP2^EGFP(△59) and BLTP2^Halo full length (FL). BLTP2^EGFP(△59) show reduced localization relative to BLTP2^Halo FL at ER-PM contact sites in U2OS cells (arrowhead) and at tubular structures in HeLaM cells.

**Figure S4.**
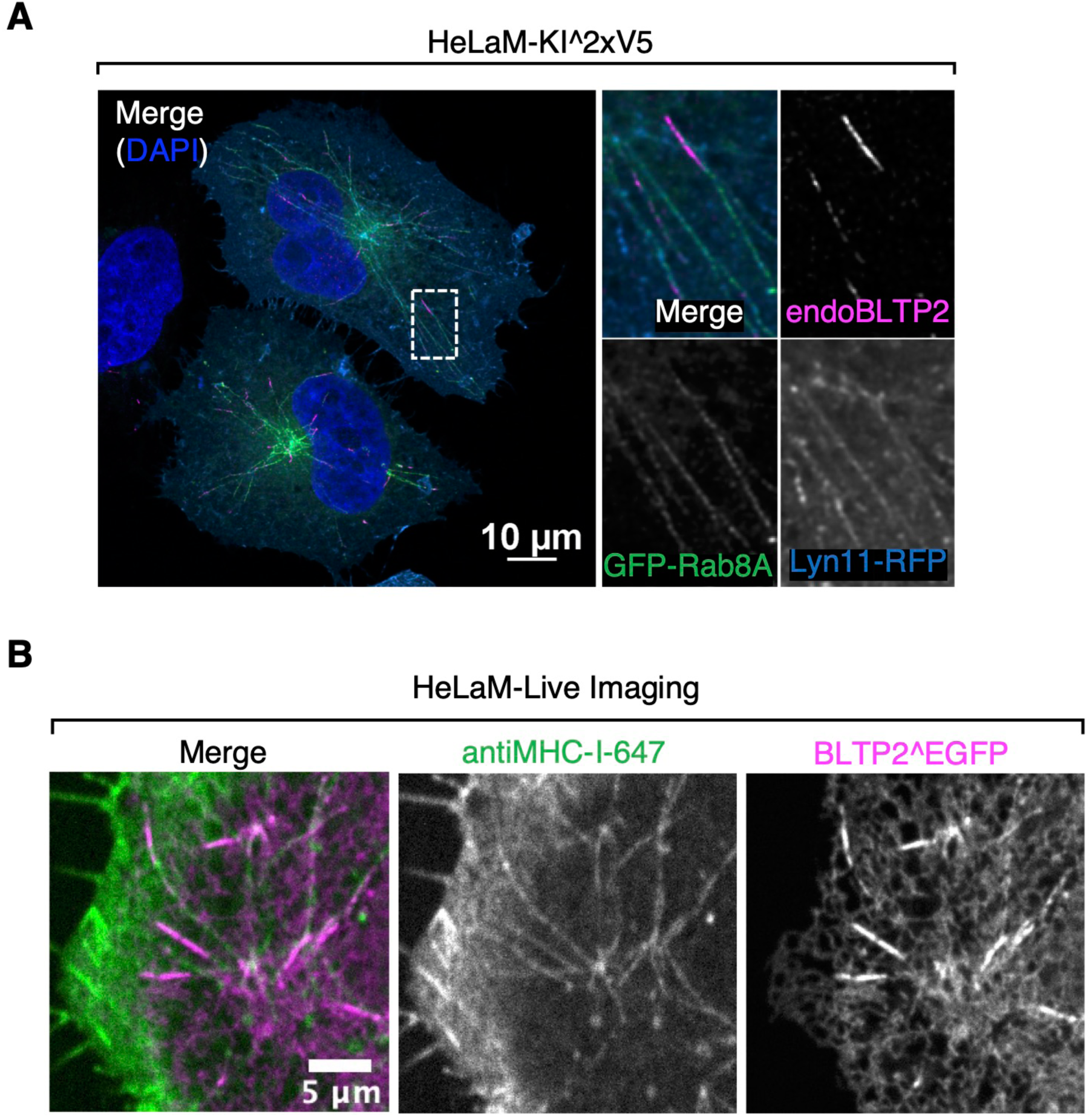
BLTP2-positive tubules have characteristics of tubular recycling endosomes. (A) BLTP2-positive tubules are positive for Rab8A. (B) A fluorescent labeled MHC-I antibody added to HeLaM cells expression BLTP^EGFP labels the tubules.

**Figure S5.**
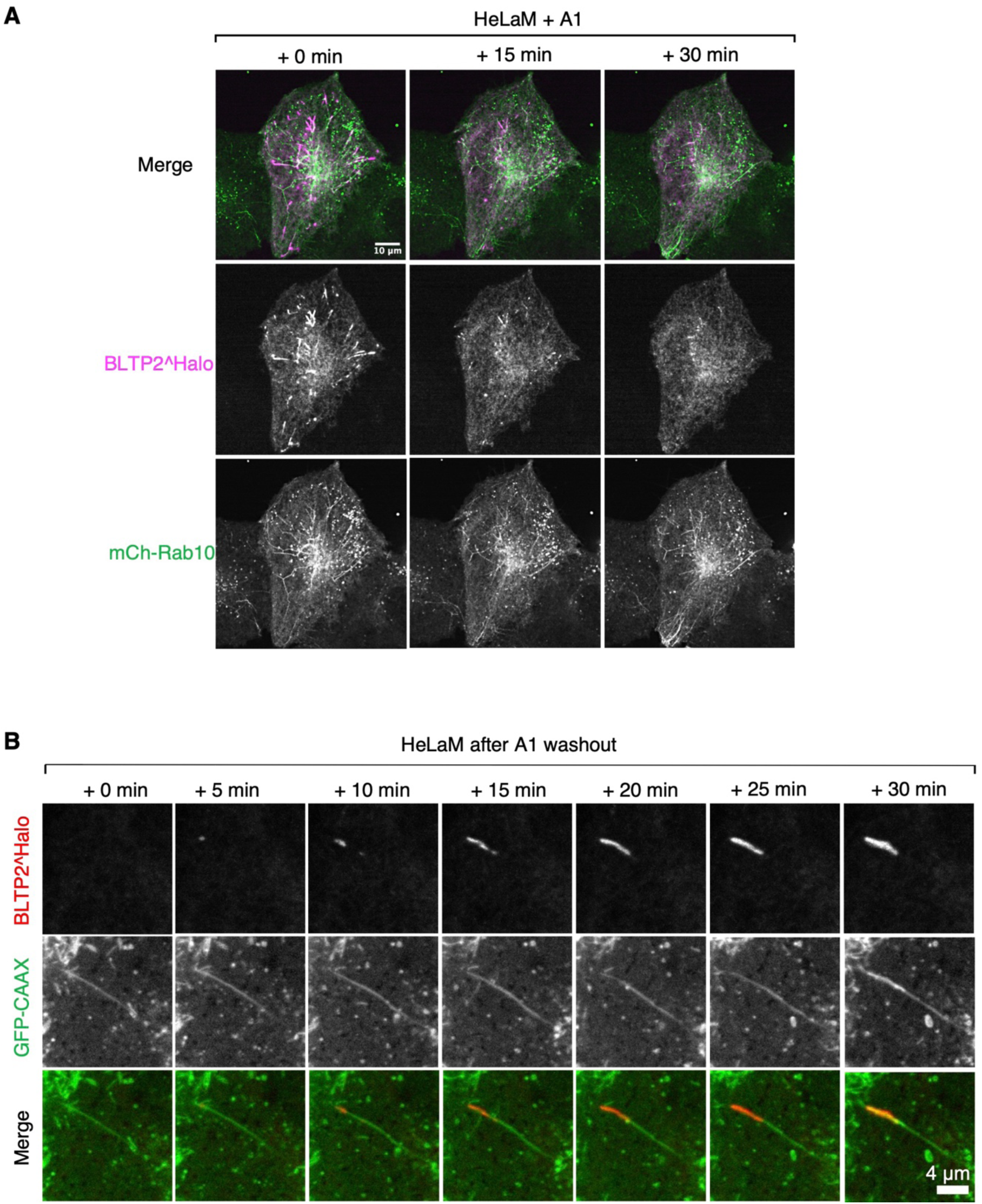
PI4P depletion does not disrupt the Rab10 tubular endosome network. (A) The disassociation of BLTP2 after A1 treatment does not correlate with a disruption of the entire Rab10-positive tubular endosome network in HeLaM cells. (B) BLTP2 re-establish contact with tubular endosomes after A1 washout.

**Figure S6.**
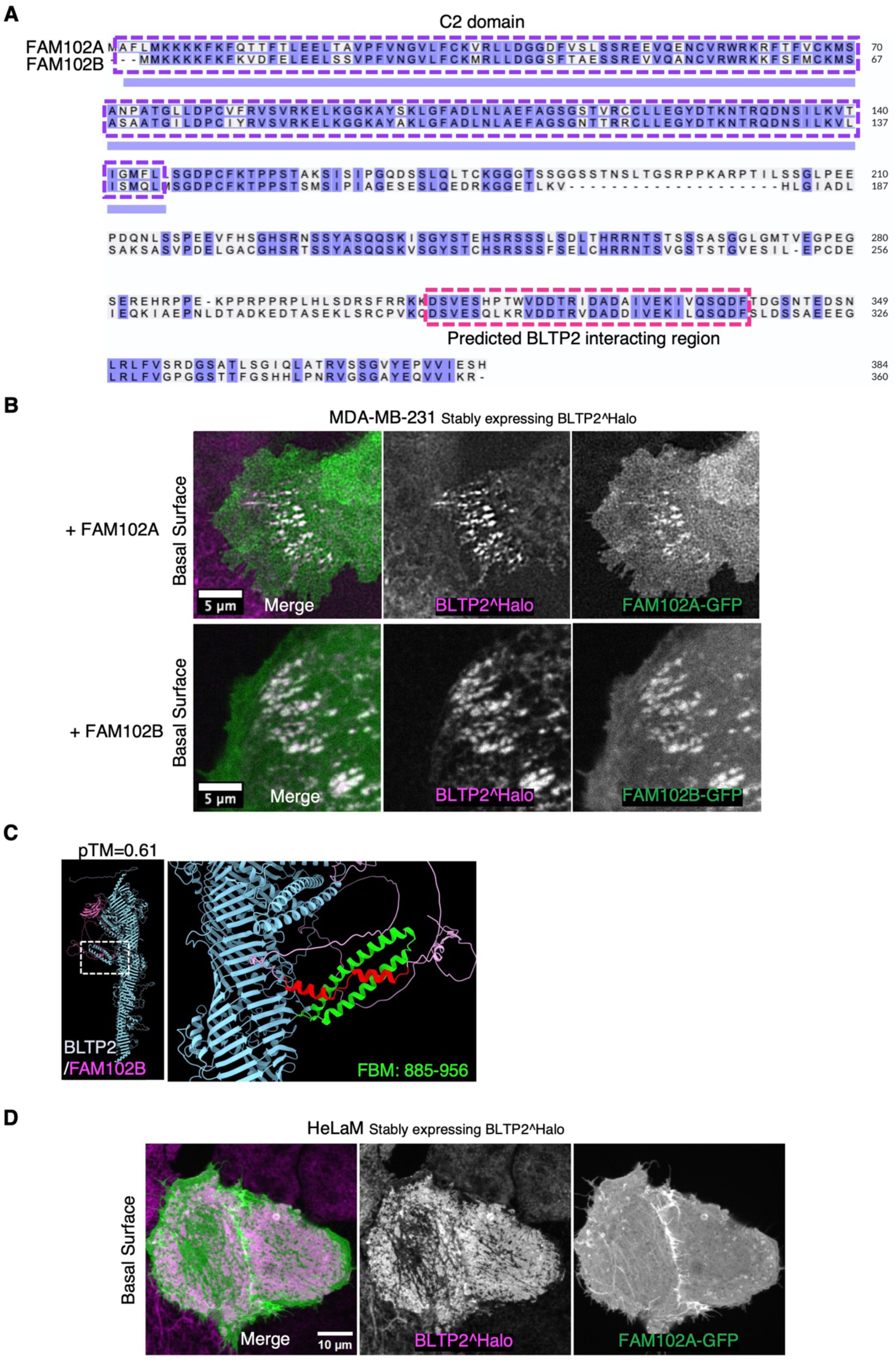
Both FAM102A and FAM102B interact with BLTP2. (A) Sequence alignment of human FAM102A and FAM102B where their N-terminal C2 domains (purple dashed box) and their C-terminal predicted BLTP2 interacting regions (red dashed box) are highlighted. (B) Co-localization of FAM102A-GFP (Top) and FAM102B-GFP (Bottom) with BLTP2^Halo at ER-PM contact sites in MDA-MB-231 cells. (C) AlphaFold3 predicted interaction between FAM102B and BLTP2. (D) FAM102A-GFP recruits BLTP2^Halo to ER-PM contact sites in HeLaM cells.

**Figure S7.**
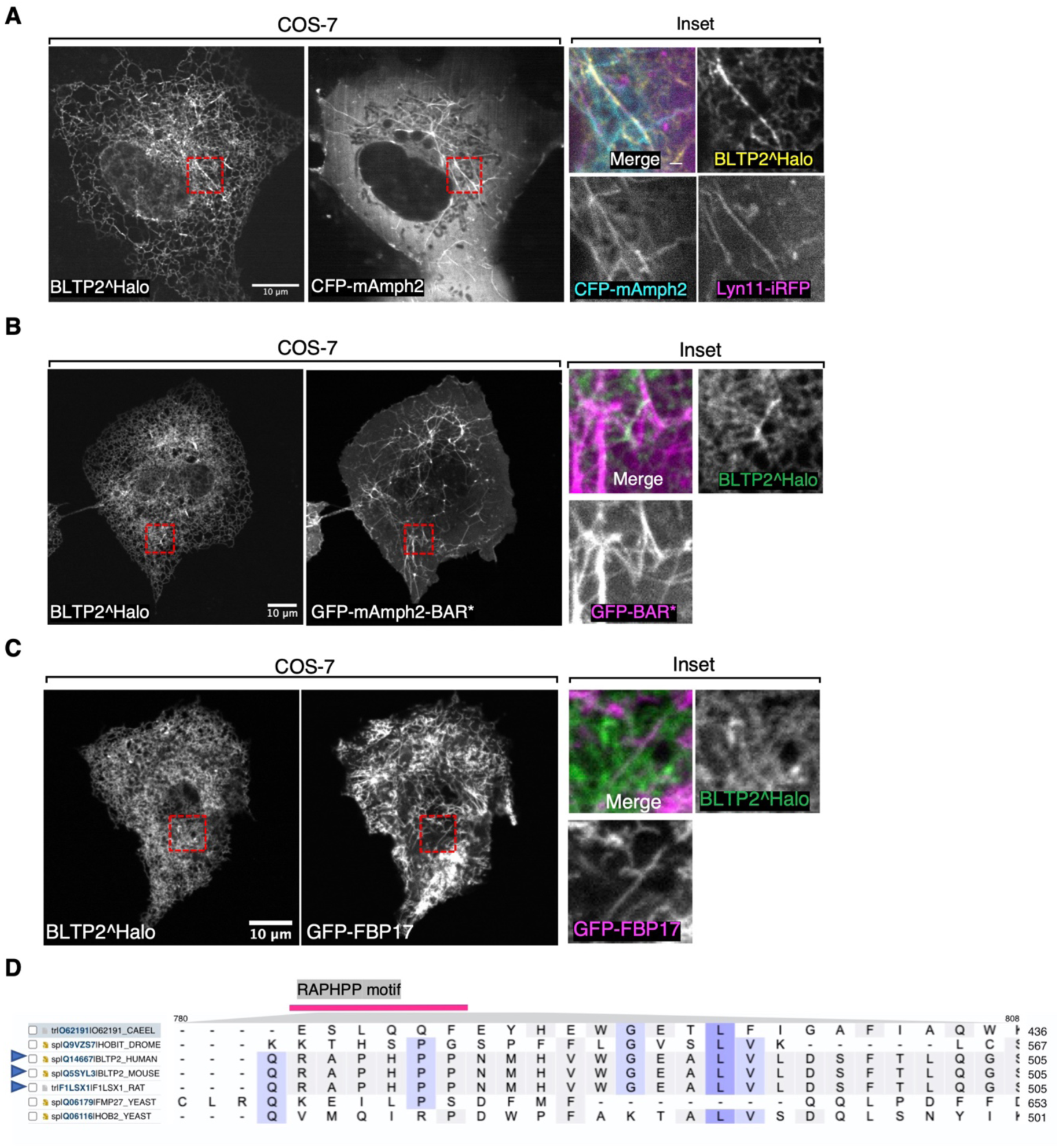
BLTP2 interacts with amphiphysin 2 in an SH3 domain dependent manner. (A) COS-7 cells showing that BLTP2^Halo is recruited to PM tubular invaginations induced by expression of CFP-mAmph2. (B) COS-7 cells showing that BLTP2^Halo is not recruited to PM tubular invaginations induced by expression of GFP-mAmph2 BAR* which lacks the SH3 domain. (C) COS-7 cells showing that BLTP2^Halo is not recruited to PM tubular invaginations induced by expression of GFP-FBP17. (D) Sequence alignment of the region comprising the RAPHPP motif among BLTP2 from several organisms. The RAPHPP motif is conserved among mammalian species (blue arrowhead).

**Figure S8.**
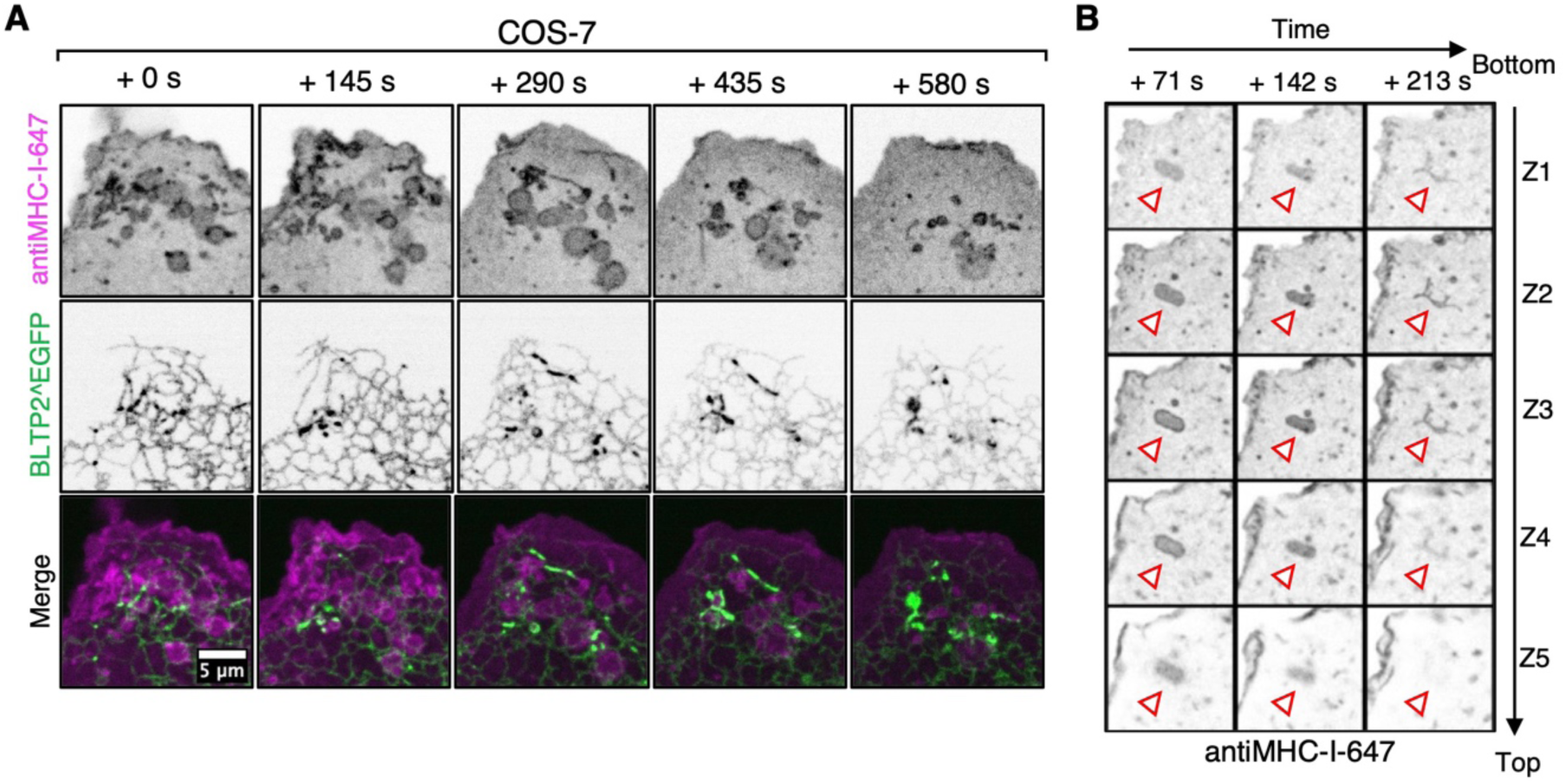
BLTP2 is recruited to macropinosomes undergoing fusion with the PM in COS-7 cells. (A) Acute recruitment of BLTP2^EGFP to macropinosomes (labeled by internalized fluorescent anti MHC-I antibodies) in the process of fusing with the PM. (B) Z-stack of the time lapse images shown in Figure 6A proving that the macropinosome is not moving out from the focal plane during the imaging session.

**Figure S9.**
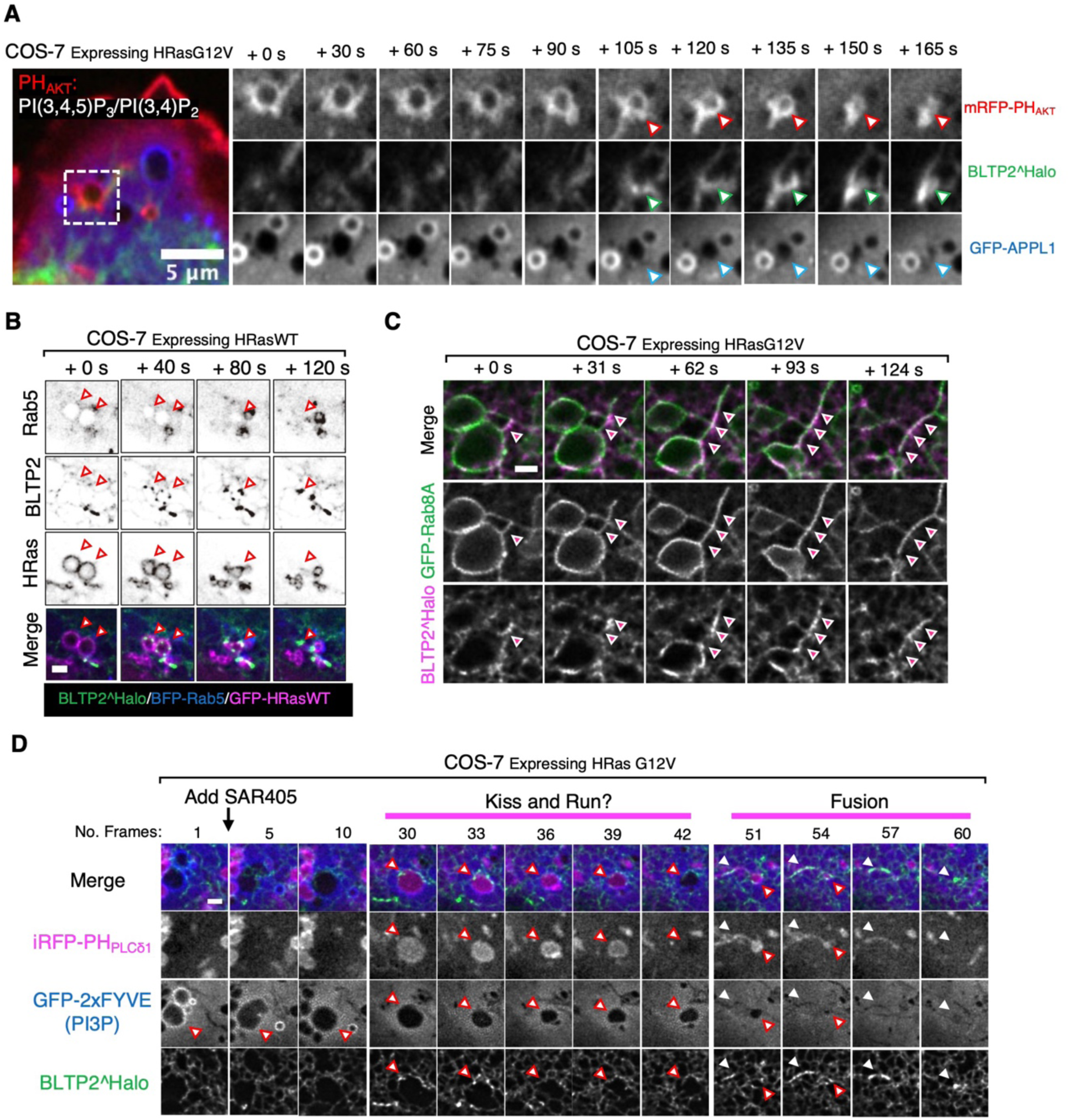
BLTP2-positive recycling macropinosomes are not matured into early endosomal stage. (A) PH_AKT_ also label nascent macropinosomes. The recycling macropinosomes are positive for PH_AKT_ while recruiting BLTP2 and are not positive for APPL1. (B) BLTP2-positive recycling macropinosomes are not positive for early macropinosome marker Rab5. (C) BLTP2-positive recycling macropinosomes are positive for exocytic factor Rab8A. (D) After treating cells with VPS34 inhibitor SAR405, the PI3P positive early macropinosomes can reverse to PI(4,5)P_2_ and recycle back to the PM.

## Supplemental Videos legends

**Video S1**. HeLaM cell expressing BLTP2^EGFP and mCherry-Rab10 showing the recovery of mCherry-Rab10-positive tubules after the wash out of nocodazole, and the recruitment of BLTP2^EGFP at their tips when they reach the PM.

**Video S2**. HeLaM cell expressing BLTP2^Halo and mCherry-Rab10 showing dissociation of BLTP2^Halo from mCherry-Rab10-positive tubules after the addition of the A1 compound. The mCherry-Rab10-positive tubule network is not disrupted.

**Video S3**. HeLaM cell expressing BLTP2^Halo and GFP-CAAX showing recovery of BLTP2^Halo-positive contacts on a GFP-CAAX-positive tubule after the wash out of the A1 compound. The wash-out is carried out after A1 treatment for one hour.

**Video S4**. COS-7 cell co-expressing BLTP2^Halo (white), FAM102A-GFP (not shown) and M1R-blank (not shown) showing disruption of BLTP2^Halo-dependent ER-PM contacts after the addition of Oxo-M, with the dispersion of BLTP2^Halo throughout the ER and the re-establishment of such contacts after the addition of atropine.

**Video S5**. COS-7 cell co-expressing BLTP2^Halo (white), FAM102B-GFP (not shown) and M1R-blank (not shown) showing disruption of BLTP2^Halo-dependent ER-PM contacts after the addition of Oxo-M, with the dispersion of BLTP2^Halo throughout the ER and the re-establishment of such contacts after the addition of atropine.

**Video S6**. COS-7 cell showing a macropinosome labeled by anti-MHC-I-647 antibodies that undergoes a drastic morphological change as it fuses with the PM. This transition correlates with the formation of BLTP2^EGFP positive ER contacts.

**Video S7**. COS-7 cell showing multiple newly formed macropinosomes labeled by antiMHC-I-647 antibodies that undergo drastic morphological change as they fuse with the PM and acquire patches of BLTP2^EGFP during this transition.

**Video S8**. COS-7 cell showing a newly formed macropinosome labeled by antiMHC-I-647 antibodies that first loses PI(4,5)P_2_ (as detected by GFP-PH_PLCδ1_) just after internalization, then re-acquires PI(4,5)P_2_ as it fuses with the PM. The acquisition of PI(4,5)P_2_ (which signals fusion with the PI(4,5)P_2_ rich PM) correlates with the formation of BLTP2^Halo positive ER contacts.

**Video S9**. COS-7 cell expressing HRas G12V and showing a newly formed PI(4,5)P_2_-positive (GFP-PH_PLCδ1_ signal) macropinosome that loses PI(4,5)P_2_, but then re-acquires it along with PM dye signal as it fuses back to the PM. BLTP2^Halo is also recruited during its fusion with the PM.

**Video S10**. BLTP2-KO HeLaM cell expressing HRas G12V showing that a subset of the internally accumulated vacuoles positive for PI(4,5)P_2_ (iRFP-PH_PLCδ1_) become labeled with the PM dye CellBrite shortly after the addition of this dye, signaling continuity of these vacuoles with the PM.

**Video S11**. BLTP2-KO HeLaM cell expressing HRas G12V, showing that upon addition to the cells of 10KD-Dextran 488 the lumen of a subset of the internally accumulated vacuoles become positive for this marker within few minutes signaling their accessibility to the extracellular medium.

